# CONSERVATION AND DIVERGENCE WITHIN THE *ARABIDOPSIS* TPL/TPR COREPRESSOR FAMILY

**DOI:** 10.64898/2026.06.02.729393

**Authors:** Benjamin L.R. Downing, Maria Pattichis, Fabian E. Vaistij, Mohamed Farawila, Edward Ghannam, Linda Nguyen, Katherine Denby, Alexander Leydon, Jennifer Nemhauser

**Affiliations:** Department of Biology, University of Washington, Seattle, WA 98195, USA; Centre for Novel Agricultural Products (CNAP), Department of Biology, University of York, York YO10 5DD, UK

## Abstract

The TOPLESS (TPL) and TOPLESS-RELATED (TPR1–TPR4) corepressors, collectively the *Arabidopsis* TPX family, are recruited by client proteins to regulate nearly every major plant regulatory program. Here, we dissect the conservation and divergence between paralogs using a combination of higher-order genetics, transcriptomics, and a synthetic repression assay. TPL, TPR1, and TPR4 were found to act as the primary repressors for many pathways, while TPR2 and TPR3 played a lesser or sometimes opposite regulatory role. Natural variation in the EAR-binding pocket subdivided the family into three subtypes: TPL/TPR1, TPR2/TPR3 and TPR4, and this variation at least partially explained observed differences between mutant phenotypes. In addition, cell-type-specific expression of EAR-containing effectors were used to tune root architecture, providing a possible route to engineering other TPX-regulated pathways. These results suggest a model where the TPX family balances robustness under stable conditions with the need for flexibility during cell fate transitions or stress responses.

## Introduction

TOPLESS (TPL) and TOPLESS-RELATED corepressors (TPR1 through TPR4 in *Arabidopsis thaliana*), hereafter referred to collectively as the TPX family, work in conjunction with diverse transcription factors to regulate expression of genes across nearly every major regulatory program in plants (Causier et al., 2012a; Lee and Golz, 2012; Liu and Karmarkar, 2008; Long et al., 2006). Indeed, TPX proteins are critical for responses to multiple hormones, flowering time control, meristem maintenance, circadian regulation, and responses to stress and other environmental cues (Chang et al., 2019; Kagale et al., 2010; Oh et al., 2014; Pauwels et al., 2010; Plant et al., 2021; Wang et al., 2013). Recruitment of clients is mediated by multiple protein-protein interaction domains (Figure 1A). The N-terminal domain, comprising Lissencephaly homologous (LisH), C-terminal to LisH (CTLH), and CT11-RanBPM (CRA) domains, binds to client transcription factors bearing short hydrophobic repression motifs, most prominently the ethylene-responsive element binding factor-associated amphiphilic repression (EAR) motif with the consensus LxLxL (Ke et al., 2015; Martin-Arevalillo et al., 2017). The C-terminal WD40 repeats form a beta-propeller that engages non-LxLxL EAR recruitment motifs including RLFGV and DLN-type sequences (Collins et al., 2019; Liu et al., 2019).

**Figure 1.**
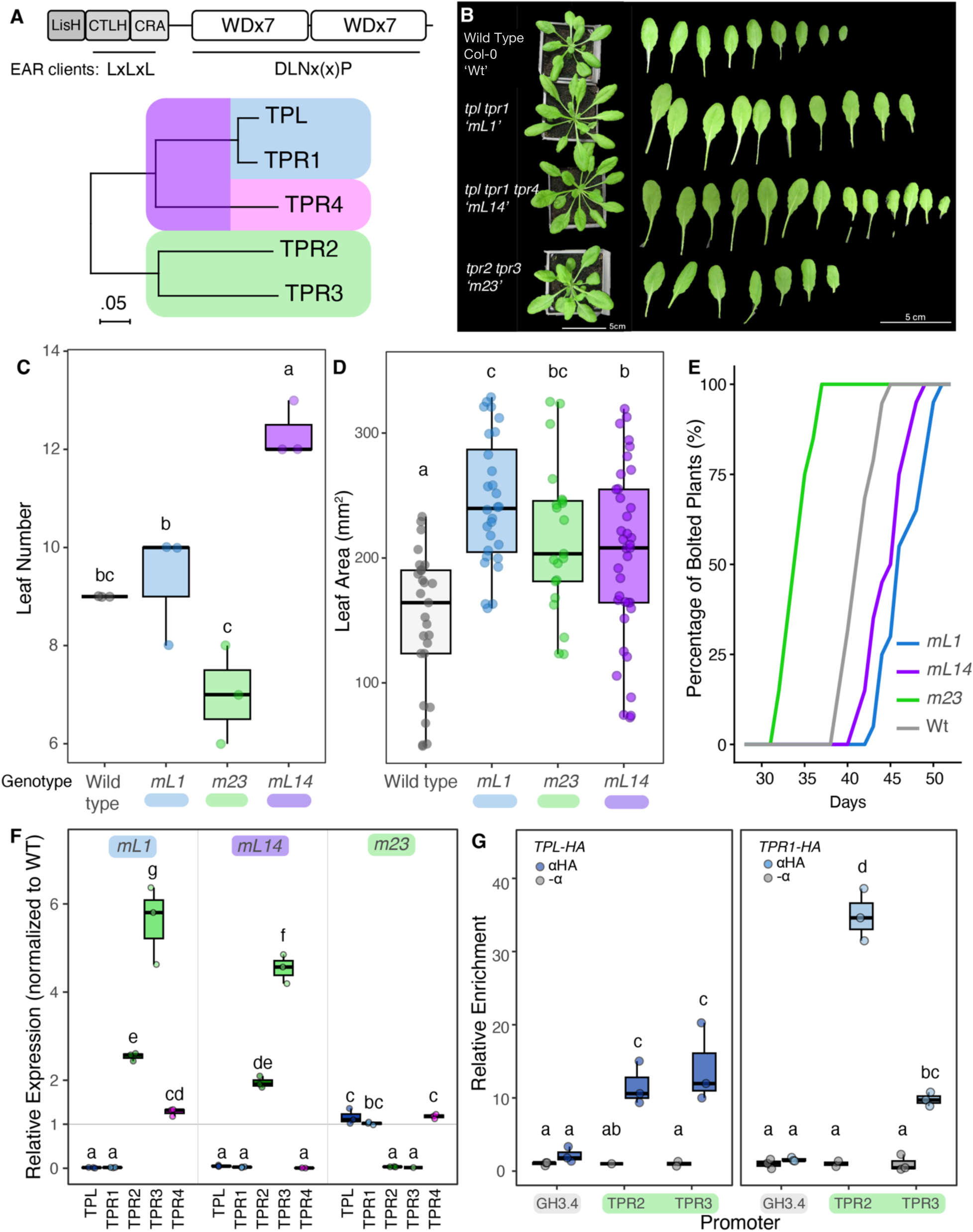
There is substantial functional redundancy within the TPX family. **(A)** Top: Schematic of conserved TPX protein domains. Bottom: Maximum likelihood phylogeny of the TPX family (JTT matrix-based model, MEGA11). Color groups used throughout all panels reflect TPX subtypes: TPL and TPR1 (blue, L1 clade), TPR4 (magenta, 4 clade), and TPR2 and TPR3 (green, 23 clade). Scale bar represents the branch length - evolutionary distance in the number of substitutions per site among the branches. **(B)** Representative rosette images and excised true leaves of Col-0 (wild type), *tpl tpr1* (*mL1*), *tpl tpr1 tpr4* (*mL14*), and *tpr2 tpr3* (*m23*) after 5 weeks of growth. **(C)** Box-and-whisker plot of rosette leaf number at 5-week old Col-0 (WT), *tpl tpr1* (*mL1*), *tpl tpr1 tpr4* (*mL14*), and *tpr2 tpr3* (*m23*) plants (3 plants per genotype). **(D)** Box-and-whisker plot of leaf area of rosette leaves of 5-week old Col-0 (WT), *tpl tpr1* (*mL1*), *tpl tpr1 tpr4* (*mL14*), and *tpr2 tpr3* (*m23*) plants (3 plants per genotype). **(E)** Line graph of days to flowering for Col-0 (WT), *tpl tpr1* (*mL1*), *tpl tpr1 tpr4* (*mL14*), and *tpr2 tpr3* (*m23*) (20 plants per genotype). Log rank test shows significant difference in bolting time p <2e^-16^. **(F)** qPCR analysis showing upregulation of *TPR2* and *TPR3* transcripts in *tpl tpr1* (*mL1*) and *tpl tpr1 tpr4* (*mL14*) backgrounds compared to wildtype. **(G)** ChIP using 2xHA-tagged TPL and TPR1 transgenic lines showing enrichment at the *TPR2* and *TPR3* promoters. Blue boxes denote binding of TPL and TPR1 to the promoters of TPR2 and TPR3 in green, with grey boxes indicating the no antibody controls. Binding to the GH3.4 promoter was performed as a negative control for enrichment. Data were analyzed by two-way ANOVA followed by Tukey’s HSD post-hoc test. Groups sharing the same letter are not significantly different (α = 0.05).

TPL, the founding member of the TPX family, was identified as a temperature-sensitive mutant (*tpl-1*) that homeotically transforms the embryonic shoot pole into a second root (Long et al., 2006, 2002). This dramatic phenotype is caused by a single amino acid substitution (N176H), proposed to inhibit function of the entire TPX family. A synthetic recapitulation of the nuclear auxin response in yeast has provided a complementary tool for dissecting TPL structure/function relationships. In the absence of auxin, Aux/IAA transcriptional repressors recruit TPL to repress expression by ARF transcription factors (Pierre-Jerome et al., 2014). The yeast synthetic system was also used to identify two autonomous repression domains within the TPL N-terminus, one in the LisH domain and one in Helix 8 of the CRA domain (Leydon et al., 2021). Helix 8 directly contacts the Mediator middle module subunits MED21 and MED10 and requires this interaction to maintain full repression (Leydon et al., 2021). Additional work revealed that the TPL LisH domain nucleates preassembly of the transcriptional preinitiation complex, including DSIF components SPT4 and SPT5, as well as the TFIID subunit TAF5 (Leydon et al., 2025). Together, these interactions support a model in which TPL (and likely other TPX family members) does not merely inhibit transcription but actively primes target loci for rapid and coordinated transcriptional bursts upon relief of repression (Leydon and Nemhauser, 2026).

Most studies to date have treated TPX proteins as largely interchangeable, reflecting the mild phenotypes of single loss-of-function mutants in each of the family members (Causier et al., 2012a; Long et al., 2006; Plant et al., 2021). Recent work in plant immunity suggests that members of the TPX family may not be fully functionally redundant. TPR1 interacts with the TIR-NBS-LRR receptor SNC1 to directly repress negative regulators of defense, including the cyclic nucleotide-gated channel genes *DND1* and *DND2*, thereby promoting immune activation (Zhu et al., 2010). In contrast, TPR2 and TPR3 were identified as negative regulators of SNC1-conditioned autoimmunity: loss of either suppressed the stunted growth and constitutive defense phenotypes caused by elevated SNC1 levels, placing them in functional opposition to TPR1 within the same pathway (Garner et al., 2021). Specific TPX family members have also been identified as susceptibility genes for *Fusarium* wilt in both tomato and *Arabidopsis* (Aalders et al., 2024). EAR-containing pathogen effectors—secreted proteins that manipulate host cellular processes to promote infection—from oomycetes (Harvey et al., 2020), fungi (Bindics et al., 2022; Darino et al., 2021; Navarrete et al., 2021), and bacteria (Gryffroy et al., 2023b) directly interact with TPX proteins and likely compete with host TPX clients. While it is likely that there are differences in affinity between a given effector and each TPX protein, this expectation remains largely untested.

Here, we combined genetic, genomic, and synthetic approaches to examine conservation and divergence within the *Arabidopsis* TPX family. We found that under standard growth conditions, the TPX family displayed substantial but asymmetric functional redundancy. TPL, TPR1, and TPR4 appeared to act as the primary repressors for many pathways, including the regulation of leaf number, flowering time, many aspects of immune responses and auxin-regulated features of root architecture. In contrast, loss of TPR2 and TPR3 resulted in minimal or opposite impacts on these same phenotypes, and both genes in this TPX subclade were found to be directly repressed by TPL and TPR1. During pathogen challenge with *Botrytis cinerea*, functional asymmetry was mirrored in sharply divergent transcriptional responses between wild type, *tpl tpr1*, *tpl tpr1 tpr4*, and *tpr2 tpr3*. In addition, synthetic assays connected sequence variation within Helix 8 in the TPR2/TPR3 clade with reduced recruitment of Aux/IAA clients, regardless of their EAR sequence. Finally, we found that LxLxL EAR-containing pathogen effectors, exemplified by the *Hyaloperonospora arabidopsidis* effector RxL21, can compete with endogenous Aux/IAA proteins for TPX binding, shifting auxin response thresholds and accelerating lateral root emergence. Together, these results support a model where divergence in TPX family member function is highly buffered until the plant is challenged by rapid state changes during development or immunity.

## Results

### Cross-regulation and functional divergence within the TPX family

The five Arabidopsis TPX proteins share an identical domain organization with a high degree of sequence conservation and low values of calculated evolutionary distance. (Figure 1A) (Causier et al., 2012b). Single loss-of-function mutants in TPX family members have mild if any phenotypes (Causier et al., 2012a; Long et al., 2006; Plant et al., 2021). To counteract the redundancy implied by previous studies, we assayed several quantitative phenotypes in mutant combinations that reflect phylogenetic relationships, namely *tpl tpr1*, *tpl tpr1 tpr4*, and *tpr2 tpr3*. We found that the number of rosette leaves of 5-week-old plants was significantly increased in *tpl tpr1 tpr4* compared to wild-type, and total leaf area was increased in all higher order mutants (Figure 1 B-D). Flowering time was accelerated in *tpr2 tpr3* when compared to wild type, a phenotype not seen in either *tpl tpr1* or *tpl tpr1 tpr4* mutants (Figure 1E). We also found cross-regulation between family members, likely contributing to the apparent functional compensation. *TPR2* and *TPR3* transcript levels sharply increased in *tpl tpr1* and *tpl tpr1 tpr4* backgrounds (Figure 1F). Chromatin immunoprecipitation (ChIP) showed enrichment of TPL and TPR1 at the *TPR2* and *TPR3* promoters, consistent with direct repression of their expression (Figure 1G).

Previous work found that the *tpl tpr1 tpr4* mutant has increased susceptibility to both the hemibiotrophic oomycete pathogen *Hyaloperonospora arabidopsidis* (Hpa) and the necrotrophic fungal pathogen *B. cinerea* (Harvey et al., 2020). Hpa utilizes well-documented effectors that target TPX family members to enhance pathogenicity (Harvey et al., 2020). *B. cinerea* also utilizes many effectors to enhance pathogenicity (Valero-Jiménez et al., 2019; Wei et al., 2024), and we found that approximately 10-12% of its documented effectors contain predicted EAR motifs (Table S1). Expanding the *B. cinerea* infection analysis to the other multiple TPX mutants, we found that *tpl tpr1* also has increased susceptibility compared to wild type, while *tpr2 tpr3* does not (Figure 2A).

**Figure 2.**
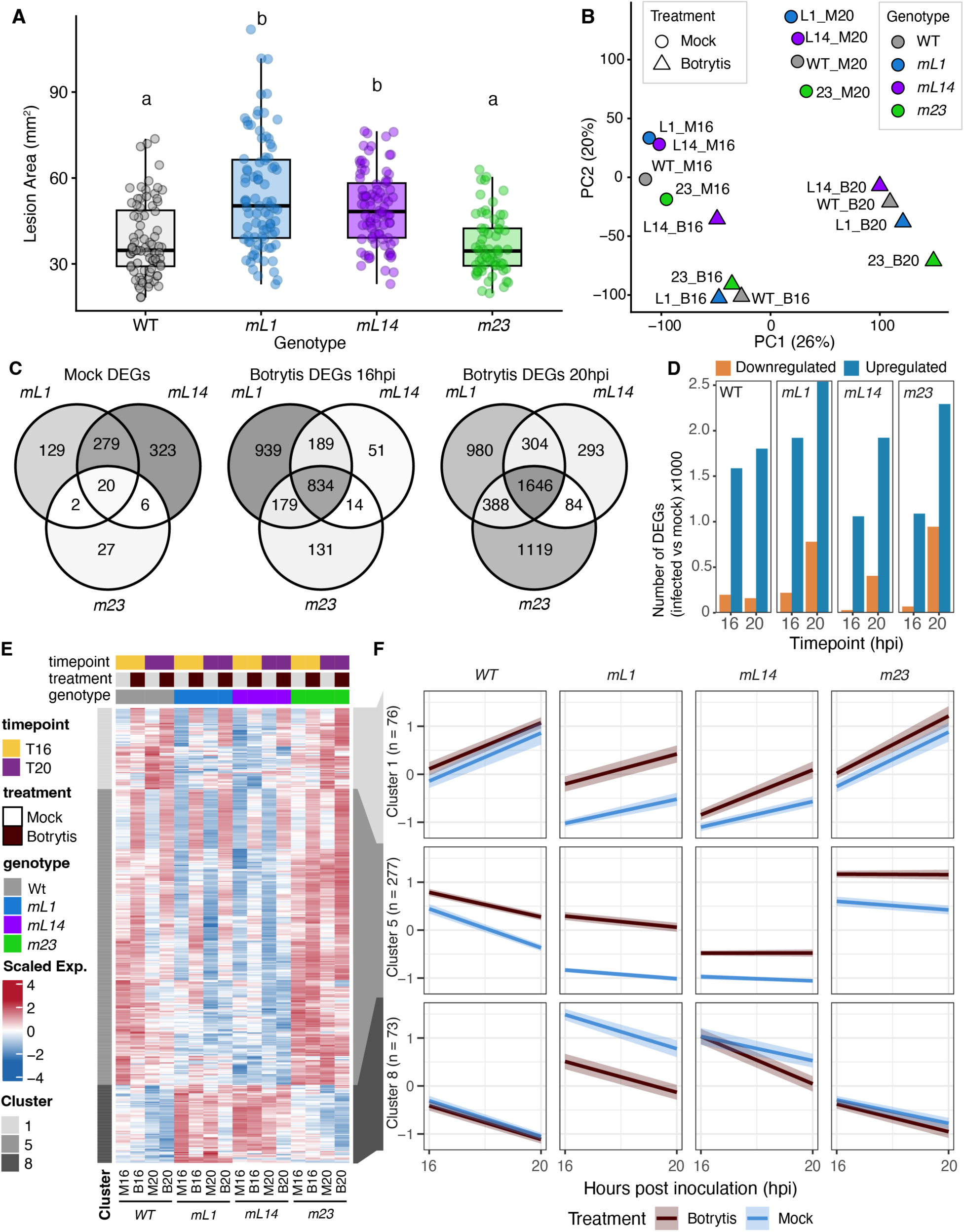
TPL/TPR corepressor mutants show altered transcriptional responses to *Botrytis cinerea* (*B. cinerea*) infection. **(A)** Quantification of *B. cinerea* lesion area on detached *Arabidopsis* leaves for wild type (WT), *tpl tpr1* (*mL1*), *tpl tpr1 tpr4* (*mL14*), and *tpr2 tpr3* (*m23*) 72 hours post inoculation (hpi), n>=63. Data were analyzed by two-way ANOVA followed by Tukey’s HSD post-hoc test. Groups sharing the same letter are not significantly different (α = 0.05). **(B)** Principal component analysis (PCA) of log2-transformed mean normalized gene expression (RNA-seq) across all detected transcripts (n = 38,676) for wild type (WT), *tpl tpr1* (*mL1*), *tpl tpr1 tpr4* (*mL14*), and *tpr2 tpr3* (*m23*) under mock and *B. cinerea* infection conditions at 16 and 20 hours post inoculation (hpi). Each point represents a single genotype × treatment × timepoint combination. Point fill indicates genotype and point shape indicates treatment (circle, mock; triangle, *B. cinerea*). Percentage of total variance explained by each principal component is indicated on the respective axis (PC1 vs PC2). **(C)** Venn diagrams showing the overlap of DEGs between *mL1*, *mL14*, and *m23* relative to WT under mock conditions, and in response to *B. cinerea* infection at 16 and 20 hpi (padj < 0.01, |log2FC| ≥ 0.5). Numbers indicate the number of DEGs unique to or shared between genotypes. **(D)** Number and direction of DEGs identified in *mL1, mL14, m23*, and WT in response to *B. cinerea* infection relative to mock (padj < 0.01, |log2FC| ≥ 0.5), grouped by timepoint and colored by direction of regulation (Upregulated, blue; Downregulated, orange). **(E)** Heatmaps showing row-scaled (z-score) normalized expression of genes belonging to selected transcriptional modules identified by hierarchical clustering of DEGs in *mL1, mL14, and m23* relative to WT under mock conditions (padj < 0.01, |log2FC| ≥ 0.5). Columns represent mean normalized counts ordered by genotype (WT, *mL1, m23, mL14*) and treatment (Mock, *B. cinerea*) at 16 and 20 hpi. Row annotations indicate cluster membership (grayscale) and whether each gene was also a DEG in response to *B. cinerea* inoculation in at least one genotype or timepoint (Upregulated, blue; Downregulated, orange; Not DE, gray). Clusters 1, 8, and 11 were selected based on interesting up or down regulation of defense genes in response to infection. **(F)** Mean scaled expression profiles (z-score) of genes within the genotype-driven modules shown in (E) across 16 and 20 hpi for each genotype under Mock (blue) and *B. cinerea* (dark red) conditions. Shaded ribbons indicate 95% confidence intervals

To understand the molecular mechanisms that underpin these phenotypic differences, we conducted transcriptomic analysis of leaves from wild type and each of the multiple mutants (*tpl tpr1*, *tpl tpr1 tpr4,* and *tpr2 tpr3*) in mock conditions and after inoculation with *B. cinerea* [16 and 20 hours post inoculation (hpi)] (Table S2). Principal component analysis (PCA) revealed similarities in gene expression among the mutant backgrounds in mock conditions, consistent with the relatively mild observed mutant phenotypes (Figure 2B). Under uninoculated conditions, the number of differentially expressed genes (DEGs) compared to wildtype in *tpl tpr1* and *tpl tpr1 tpr4* mutants was much higher than the *tpr2 tpr3* mutant (Figure 2C, Table S2). Hierarchical clustering was conducted on the set of DEGs from all the mutants compared to wild type (under mock conditions) and GO-term enrichment was performed on the clusters (Figure 2 – figure supplement 2). Expression of jasmonic acid (JA) signaling and JA response genes, along with genes involved in glucosinolate biosynthesis and defense to insects, were elevated in *tpl tpr1* and *tpl tpr1 tpr4* mutants (Figure 2E, cluster 7 and 8), while salicylic acid (SA) response genes and genes involved in defense against bacterial pathogens have lower expression compared to wild type and *tpr2 tpr3* mutants (Figure 2E, clusters 1 and 5).

Pathogen challenge revealed additional areas of conservation and divergence among the paralogs. Consistent with partial redundancy between family members, many DEGs (*B. cinerea* v. mock inoculated) are shared between all genotypes, including wildtype, (Figure 2C-D; Figure 2 – figure supplement 3, Table S2). The vast majority of these shared DEGs are upregulated in response to pathogen infection, with almost no overlap in downregulated genes (Figure 2 – figure supplement 3), and these common upregulated genes are enriched for GO terms such as responses to wounding and pathogens (Figure 2 – figure supplement 4).

Looking at genotype-specific responses to *B. cinerea*, upregulated genes specific to each genotype are enriched for similar functions in defense (Figure 2- figure supplement 4). However, amongst downregulated genes each genotype-specific gene set is enriched for different functions: *tpr2 tpr3*-specific genes were enriched in GO terms associated with response to light intensity and photosynthesis, contrasting with *tpl tpr1*-specific downregulated DEGs, which were associated with glucosinolate and starch metabolism, and *tpl tpr1 tpr4*-specific downregulated DEGs which showed enrichment for auxin signaling and response pathways (Figure 2- figure supplement 5).

### Sequence variation in Helix 8 determines efficiency of client recruitment

Alignments of the TPX family revealed a patch of natural sequence variation in Helix 8 of the CRA domain. This region is positioned within the EAR-binding pocket, which is the interface through which TPX proteins engage the LxLxL motif on Aux/IAA repressors and other client proteins (Figure 3A). Phylogenetic analysis of Helix 8 alone recapitulated the same three-subtype organization seen in the phylogeny based on full length sequences: TPL/TPR1 (L1), TPR4 (4) and TPR2/TPR3 (23) (Figure 3B). To directly test whether Helix 8 variation affects repression function, we leveraged our synthetic auxin response system in yeast (*At*ARC*^Sc^*) where TPLN188 (the first 188 amino acids of TPL, encompassing the LisH-CTLH-CRA domains) is sufficient for full repression of a reporter (Leydon et al., 2021) (Figure 3C). To make it possible to distinguish effects on repression itself from effects on recruitment of a client, we implemented two versions of *At*ARC*^Sc^*. In the first configuration, referred to here as fused, the client IAA3 protein was expressed as a fusion protein with TPLN188 carrying each Helix 8 variant (i.e., the original TPL/TPR1 sequence [H8-L1] or the TPR2/TPR3 [H8-23] or TPR4 sequence [H8-4] swapped into the TPL backbone). In a second configuration, referred to here as free, IAA3 and each of the corepressor variants were expressed as separate proteins. Flow cytometry revealed that H8-L1, H8-23, and H8-4 subtypes were able to repress transcription to comparable levels when fused to the client protein (Figure 3D,E). Consistent with this result, cytoplasmic split-ubiquitin (cytoSUS) assays (Asseck and Grefen, 2018) revealed that each TPLN188 Helix 8 subtype showed similarly strong interactions with TPLN188, IAA3, and MED21 (Figure 3 – figure supplement 3).

**Figure 3.**
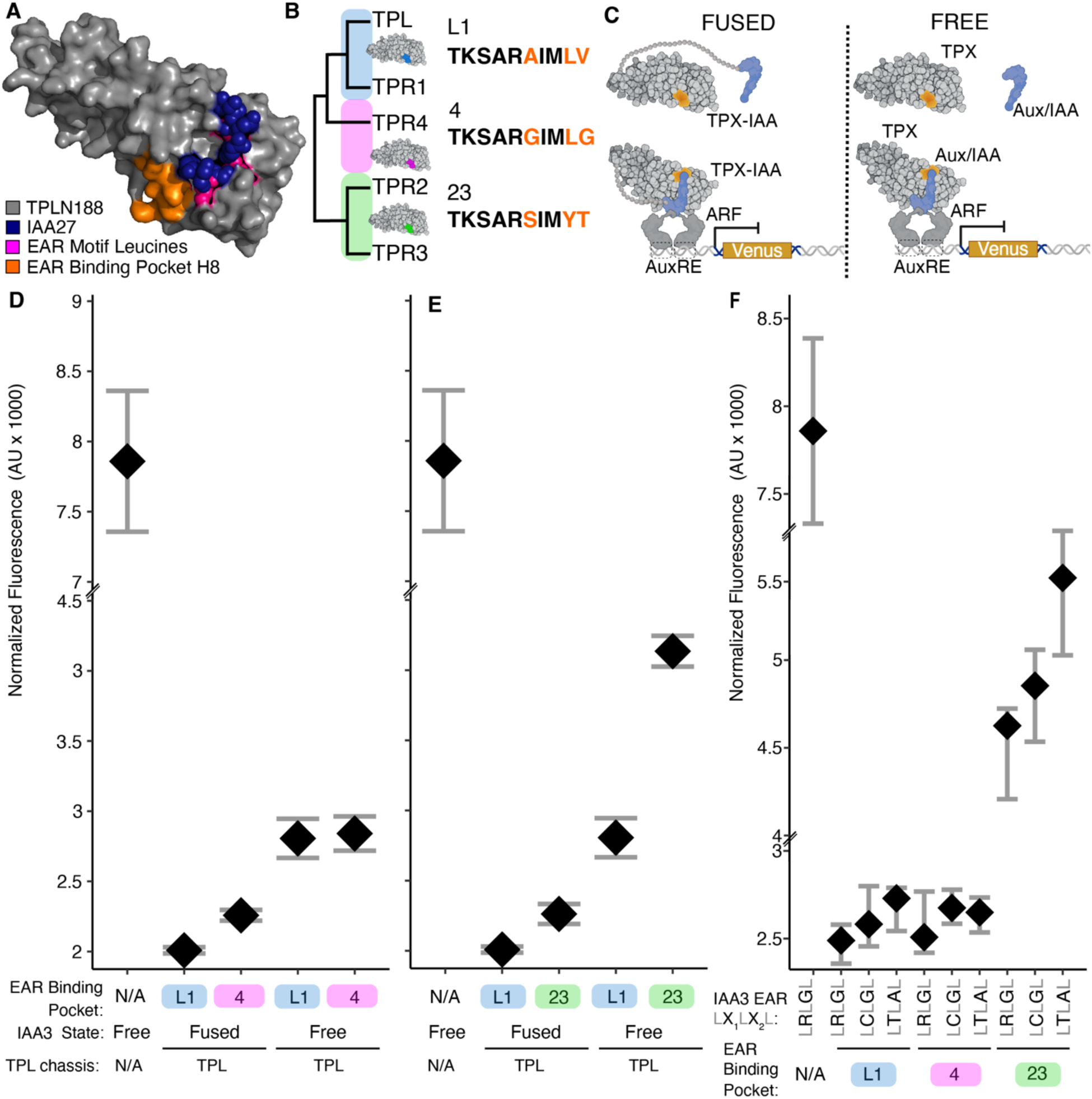
Helix 8 variation alters client recruitment efficiency. **(A)** EAR-binding pocket schematic with variant residues of Helix 8 highlighted in orange. **(B)** Phylogeny showing the unique Helix 8 sequence of each clade: L1 (TPL/TPR1, blue), 23 (TPR2/TPR3, green), 4 (TPR4, pink). **(C)** *At*ARC*^Sc^* assay schematic in fused (recruitment-bypassed) and unfused (recruitment-dependent) configurations. In the fused configuration, the TPL N-terminal domain with the Helix 8 variant of interest is directly fused to IAA3. In the unfused configuration, the corepressor and IAA3 are expressed as separate proteins. **(D-E)** mVenus reporter output normalized to cell size for Helix 8 subtypes, (D) in EAR binding pocket subtypes TPR4 and TPL/TPR1, (E) in EAR binding pocket subtypes TPR2/TPR3 and TPL/TPR1. In each case the mutations are made in TPL protein chassis. The same L1 data is plotted on each graph for comparison. **(F)** Multiple Aux/IAA LxLxL sequences confirm that differences arise from TPX-side variation in the EAR-binding pocket rather than client-side diversity. TPL chassis consists of the truncated N188 confirmation of *At*TPL. For all flow cytometry experiments, each point represents the average fluorescence of 5,000–10,000 individually measured yeast cells (a.u., arbitrary units). At least two independent experiments are included for each construct, displayed with 95% confidence intervals.

In the unfused configuration, the H8-23 subtype showed a marked reduction in repression ability relative to H8-L1 and H8-4 (Figure 3D,E). We reasoned that the difference in repression strength seen in the H8-23 subtype could reflect a divergence in affinity for distinct EAR sequences. To test this, we swapped the IAA3 LRLGL motif with the EAR from IAA14 (LCLGL) and from IAA15 (LTLAL). These specific variants were selected to reflect known differences in Aux/IAA repression activity based on EAR sequences (Cho et al., 2024); however, the effect of variation in EAR sequences had little impact on the strength of repression in our assay (Figure 3F). These results suggest that differences in Aux/IAA repression strength are not likely a result of a change in preference for binding EAR motifs among TPX paralogs.

### Pathogen effectors can modulate auxin responses

Several pathogen effectors carry canonical LxLxL EAR motifs that engage the same binding interface on TPX proteins as endogenous clients like Aux/IAAs (Figure 4A). If the H8-23 subtype recruited clients less efficiently, we predicted that it would be more susceptible to competitive displacement by an EAR-containing effector like RxL21 from Hpa (Harvey et al., 2020). In the presence of an EAR-deleted RxL21 (RxL21ΔEAR), repression in our *At*ARC^Sc^ system was maintained by all three Helix 8 subtypes, confirming that the competition occurs specifically through the LxLxL interface. In contrast, addition of full-length RxL21 (EAR: LMLTL) increased reporter expression across all Helix 8 backgrounds, consistent with competitive displacement of IAA3 from the EAR-binding pocket (Figure 4B). As predicted, the H8-23 subtype showed the largest increase in reporter expression upon RxL21 addition, consistent with its weaker recruitment of clients (Figure 4B), and sensitized the system to auxin (Figure 4 – figure supplement 2).

**Figure 4.**
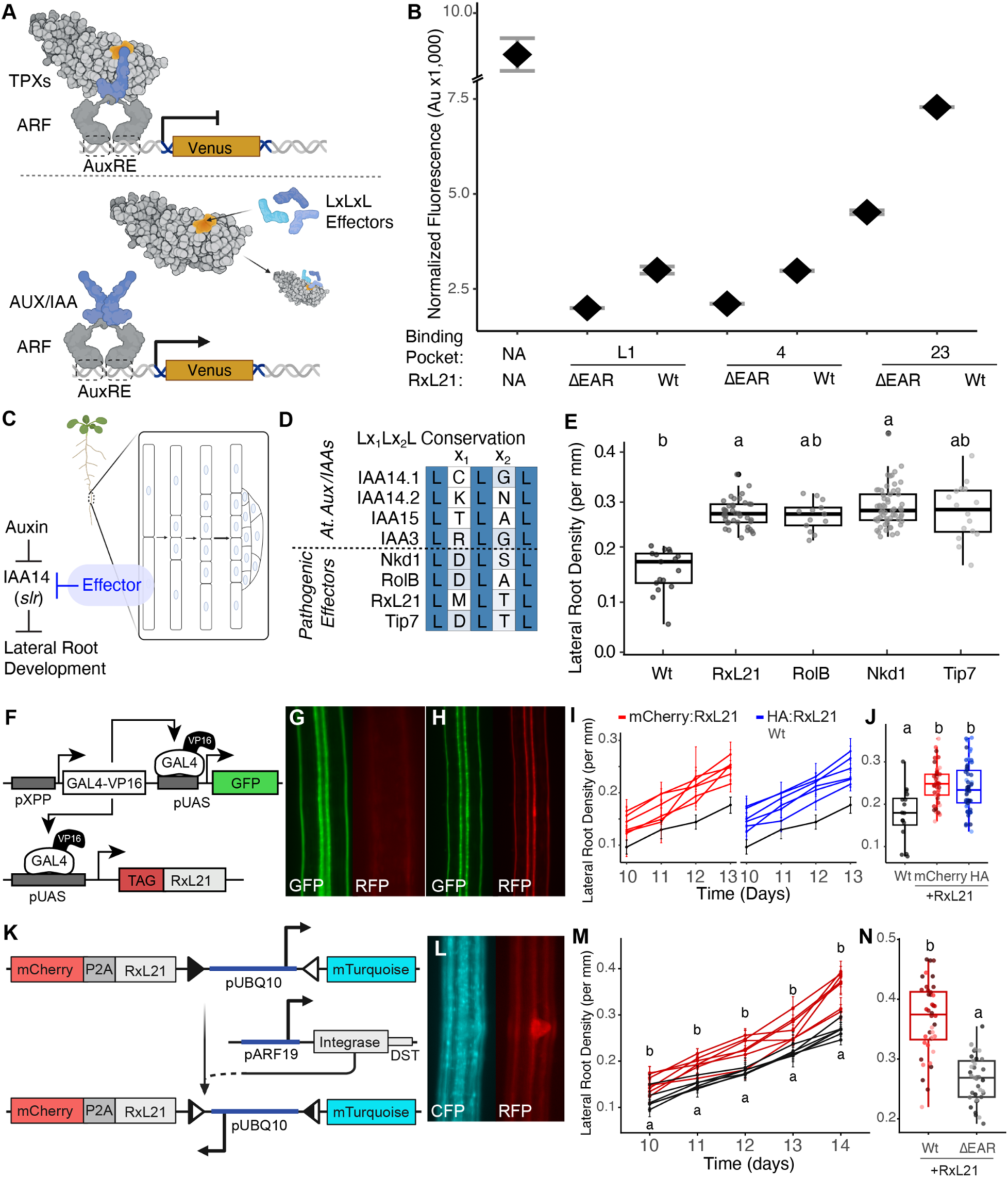
EAR-containing effectors modulate auxin sensitivity. **(A)** Competition assay schematic with RxL21 in unfused *At*ARCSc. RxL21 carries an LMLTL EAR motif and competes with IAA3 for binding to the TPX EAR-binding pocket. **(B)** RxL21 increases reporter output in all Helix 8 backgrounds (L1, 23, and 4 clades), with the strongest effect in the 23 background. Deletion of the RxL21 EAR motif abolishes the effect. For all flow cytometry experiments, each point represents the average fluorescence of 5,000–10,000 individually measured yeast cells (a.u., arbitrary units). At least two independent experiments are included for each construct, displayed with 95% confidence intervals. **(C)** Prediction of effector action on lateral root initiation. Auxin targets IAA14/S*OLITARY-ROOT* (*SLR*) for degradation which directly results in the initiation of a new lateral root. (**D**) Alignment of LxLxL EAR domains from Aux/IAA proteins and pathogenic effector proteins. **(E)** pAHP6:Effector transgenic plants show increased lateral root density relative to Col-0. **(F)** Genetic schematic of GAL4-VP16 XPP enhancer (J0121 enhancer trap line) driving tagged RxL21 expression in xylem pole pericycle cells. **(G-H)** Tagged RxL21 constructs confirm expression specificity in XPP cells. **(I)** Time-course analysis of lateral root density (days 10–13 post germination) with each line representing T2 seedlings from an independent primary transformant line. Red lines - mCherry:RxL21, blue lines - HA:RxL21, black line – parental UAS driver line control. **(J)** Lateral root density of individual lines at day 13. Boxes represent the interquartile range with median; points represent individual seedlings. In both panels, letters indicate statistically significant differences between WT, HA:RxL21, and mCherry:RxL21 groups as determined by pairwise comparison of estimated marginal means with Sidak adjustment (α = 0.05). **(K-L)** iInducer construct schematic: pUBQ10-driven mTurquoise is inverted by PhiC31 integrase (pARF19) to switch to mCherry-P2A-RxL21. **(H)** pARF19-triggered pUBQ10::mCherry-P2A-RxL21 iInducer expression in all lineages derived from pARF19 expression in XPP cells. Root microscopy images showing integrase switching of the iInducer construct in early developing lateral roots. **(M-N)**. Lateral root density (LR/mm) of WT (red) and RxL21-ΔEAR (black) iInducer lines measured over a five-day time-course (days 10–14 post-germination) and at day 14 endpoint. Each individual line represents T2 seedlings from independent insertion lines. n = 25–40 plants per line with each line representing a single seedling. Error bars indicate ± SEM. **(N)** Lateral root density of individual lines at day 14. Boxes represent the interquartile range with median; points represent individual plants. In both panels, letters indicate statistically significant differences between WT and RxL21-ΔEAR groups as determined by pairwise comparison of estimated marginal means with Sidak adjustment (α = 0.05).

To test whether effector competition for clients could work similarly in the far more complex environment of a plant cell, we expressed EAR-containing effectors in xylem pole pericycle (XPP) cells, the cell type in which auxin-induced degradation of IAA14 triggers initiation of a new root (Figure 4C). If EAR-containing effectors displace endogenous Aux/IAA repressors from TPX proteins, we reasoned that expression in XPP cells should mimic the auxin response thereby promoting lateral root formation. We also tested whether variation among EAR domains (Figure 4D) would impact the degree of dysregulation of the auxin response. Specifically, we used pAHP6 to drive expression of effectors from Hpa (RxL21), rhizogenic *Agrobacterium* (RolB), and the fungus *Ustilago maydis* (Nkd1 and Tip7) (Gryffroy et al., 2023a; Khan et al., 2024; Moreira et al., 2013; Navarrete et al., 2022a). In all cases, we found a trend of increased average lateral root density with expression of these natural TPX-sponges, with significant changes found in lines expressing RxL21 and Nkd1 (Figure 4E). Given the large number of effectors injected during most infections which can have diverse molecular targets, it was notable that expression of a single effector could lower the threshold for auxin response.

To visualize the spatial pattern of effector expression, we used the J0121 enhancer trap line, which drives GAL4-VP16 expression specifically in XPP cells (Figure 4F). Tagged RxL21 constructs confirmed expression specificity and reproducibility in this system through visualization of mScarlet reporter expression colocalized with GFP in XPP cells (Figure 4G, H). Time-course analyses demonstrated accelerated lateral root emergence between 10 and 13 days after germination and significantly increased lateral root density after 13 days (Figure 4I-J). The same pattern was observed when a serine-integrase-based recombination system was used to drive higher expression of RxL21 (and mScarlet) from a strong pUBQ10 promoter whose activity was restricted to lateral root initial cells (Figure 4K). In this approach, transgenes are selectively switched from driving expression of mTurquoise to expression of mScarlet-P2A-RxL21 (Figure 4K-L) (Guiziou et al., 2023). The resulting transgenic lines had increased lateral root density and total lateral root number relative to controls (Figure 4M-N).

### Auxin responses in roots and flowers are primarily mediated by TPL, TPR1, and TPR4

As RxL21 was able to alter root architecture by effectively lowering the dosage of available TPX proteins, we predicted that root development might be particularly sensitive to loss of TPX function. We tested this prediction by examining developmental phenotypes in higher-order mutant combinations, focusing on loss of function of *TPL*, *TPR1* and *TPR4* as these were the most effective repressors in our synthetic auxin response assays. Primary roots of *tpr1*, *tpl tpr1* and *tpl tpr1 tpr4* had a similar overall morphology as wild-type plants (Figure 5A-C). A dose response experiment revealed that *tpl tpr1 tpr4* showed significant auxin hypersensitivity (p = 0.018) (Figure 5D). We next tested all genotypes with a single auxin concentration (6.5 nM) that appeared to maximally differentiate responses, and found that both *tpl1 tpr1* and *tpl tpr1 tpr4* were significantly more sensitive than wild-type (Figure 5 - figure supplement 2). This pattern supported the conclusion that *TPL*, *TPR1*, and *TPR4* together buffer auxin response thresholds. Consistent with this interpretation, the length of root hairs, known to be auxin regulated (Velasquez et al., 2016), was modestly increased in *tpr1* and *tpl tpr1* mutants, but substantially increased in *tpl tpr1 tpr4* mutants (Figure 5E-F). Auxin hypersensitivity phenotypes were also observed in the flower, where petal number became less canalized as TPX dosage decreased. The *tpl tpr1 tpr4* triple mutant shifted towards an average of five or six petals compared with the four petals seen in essentially every wild-type flower (Figure 5 - figure supplement 4).

**Figure 5.**
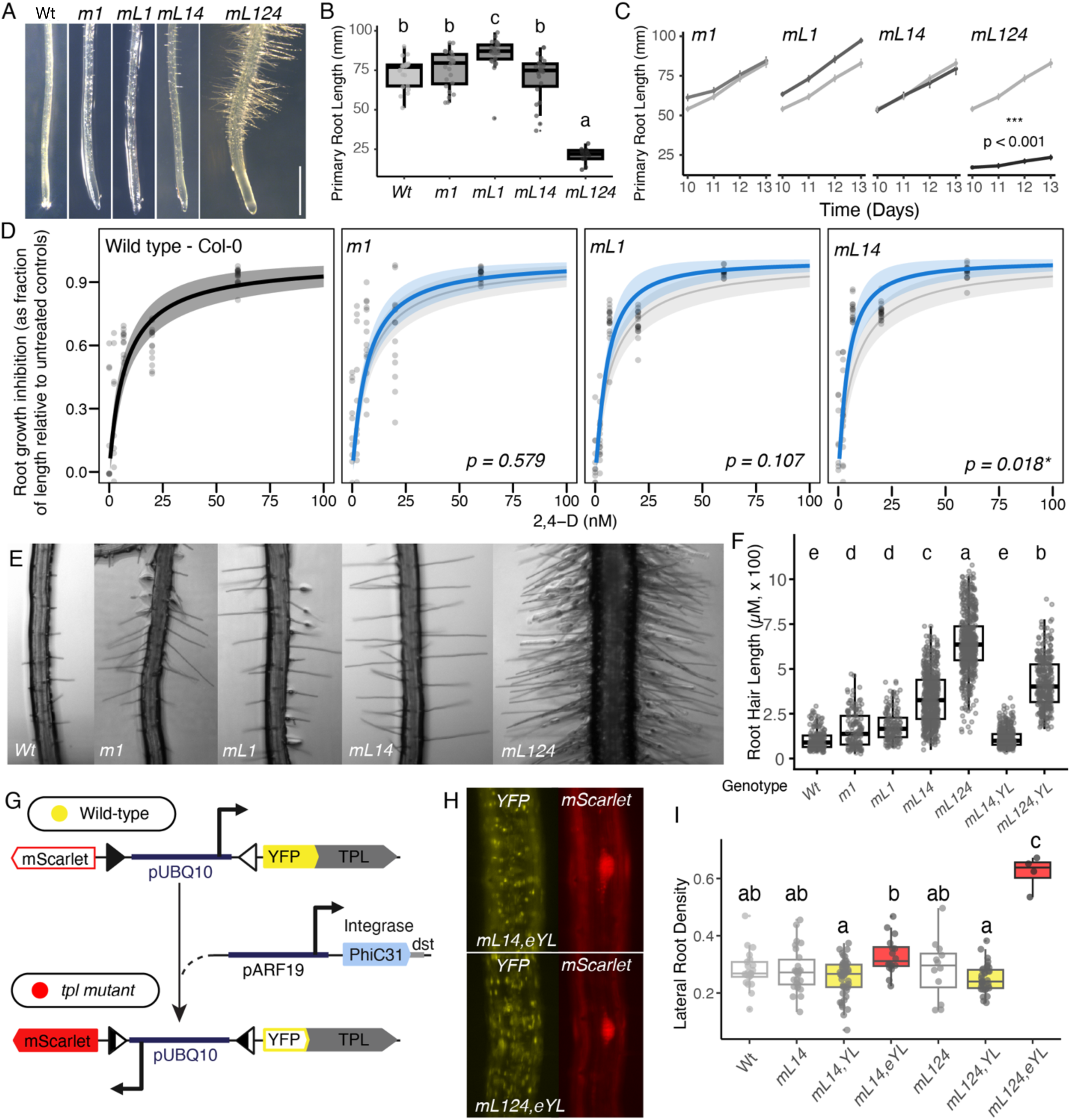
Loss of the dominant TPX clade causes auxin hypersensitivity and developmental disruption. **(A)** Dissection microscope images of root tips from Col-0 (WT), *tpr1* (*m1*), *tpl tpr1* (*mL1*), and *tpl tpr1 tpr4* (*mL14*). **(B-C)** Primary root lengths for each Col-0, *tpr1* (*m1*), *tpl tpr1* (*mL1*), and *tpl tpr1 tpr4* (*mL14*) were measured over a time-course from 10-13 days post germination (3 replicates of 8 seedlings per genotype). **(B)** Root lengths quantified at Day 13. Data were analyzed by two-way ANOVA followed by Tukey’s HSD post-hoc test. Groups sharing the same letter are not significantly different (α = 0.05). **(C)** Primary root lengths graphed over time, slopes were compared across genotypes using pairwise contrasts of estimated marginal trends (emtrends) from a linear model including a day × genotype interaction, with Tukey’s HSD adjustment for multiple comparisons. **(D)** Root growth inhibition across 2,4-D concentrations for Col-0, *tpr1* (*m1*), *tpl tpr1* (*mL1*), and *tpl tpr1 tpr4* (*mL14*). Black curves show fitted dose-response models with 95% CI (gray). Only *mL14* shows significant auxin hypersensitivity (p = 0.018). **(E)** DIC micrographs of root hair zones showing progressive enhancement of root hair density and length from Col-0 through *tpl tpr1 tpr2 tpr4* (*mL124*). **(F)** Average root hair length increases with progressive depletion of TPX proteins (n=2203 root hairs). Data were analyzed by two-way ANOVA followed by Tukey’s HSD post-hoc test. Groups sharing the same letter are not significantly different (α = 0.05). **(G)** iEraser conditional complementation construct schematic: pUBQ10-driven YFP-TPL is inverted by PhiC31 integrase (pARF19) to switch to mScarlet. **(H)** Root microscopy images showing integrase switching of the iEraser (YL) in *tpl tpr1 tpr4* (*mL14*) and *tpl tpr1 tpr2 tpr4* (*mL124*) backgrounds, average root hair length graphed in (F). **(I)** Lateral root density per mm across genotypes including *mL14* and *mL124* mutants, corresponding YFP-TPL rescue lines (*mL14*,YL and *mL124*,YL, yellow), and integrase eraser lines (*mL14*,eYL and *mL124*,eYL, red). Data were analyzed by two-way ANOVA followed by Tukey’s HSD post-hoc test. Groups sharing the same letter are not significantly different (α = 0.05).

Based on the *At*ARC*^Sc^* results, we expected that further loss of either *TPR2* or *TPR3* in the triple *tpl tpr1 tpr4* background would result in a dramatic increase in the severity of developmental defects. To test this prediction, we generated a CRISPR/Cas9 knock-out of *TPR2* in the *tpl tpr1 tpr4* background as *TPR2* and *TPR4* are tightly linked (Figure 1 - figure supplement 1D,E). Consistent with a model where a single functioning TPX family member would drop the plant below a critical threshold of TPX dosage, the *tpl tpr1 tpr2 tpr4* mutant showed qualitative changes in root morphology and a sharply reduced growth rate (Figure 5A-C). Treatment with the auxin antagonist auxinole, a competitive inhibitor of the TIR1/AFB auxin co-receptors, was able to partially rescue this phenotype (Figure 5 - figure supplement 2). The length of root hairs on quadruple mutants was more than 6X what was observed in wild-type seedlings, and the plants exhibited increased number and density of root hairs as well as having an increased overall root diameter (Figure 5E-F). Similar trends were observed in the flower where the emergence of an *apetala2* phenotype made it impossible to quantify the impact on petal number (Figure 5 - figure supplement 4D-G,J-K). This extreme phenotype is consistent with previous studies connecting TPL with *APETALA2* repression of *AGAMOUS* (Krogan et al., 2012), and suggests that the TPX dosage needed to maintain wild-type function may vary among client proteins.

Given the central role of auxin in regulating lateral root initiation, we were surprised to find no significant change in lateral root density in any of the mutants (Figure 5I). We hypothesized that this might reflect the regulatory compensation we had observed within the family (Figure 1F), specifically that up-regulation of TPR2 and TPR3 provided sufficient TPX function to maintain repression at auxin-regulated genes in pericycle cells. To test this hypothesis, we designed an experiment using iEraser technology (Watson et al., 2024) to induce acute loss of TPL function at the moment of lateral root initiation, thereby limiting the time for potential compensation to take effect. In these iEraser experiments, *tpl tpr1 tpr4* and *tpl tpr1 tpr2 tpr4* mutants were transformed with two constructs: (a) a target which can switch between constitutive expression of YFP-TPL to mScarlet by inverting the direction of the promoter via serine-integrase-directed recombination and (b) a driver that directs the expression of the PhiC31 serine integrase in cells where the pARF19 promoter is active (Figure 5G). Fluorescence imaging confirmed efficient integrase-driven recombination in our transgenic lines (Figure 5H). In both the triple and quadruple mutant backgrounds, constitutive expression of YFP-TPL resulted in a slight and statistically insignificant decrease in median lateral root density compared with wild-type plants; however, in combination with the acute erasure of YFP-TPL in lateral root initial cells, this effect made it possible to detect a modest yet significant increase in lateral root density in the triple mutant (Figure 5H). The sensitization to loss of TPL function was even more obvious with the dramatic increase in lateral roots in the quadruple mutant with the iEraser constructs. These results highlight the role of cross-regulation within the family as necessary to maintain critical thresholds of TPX function during periods of cell state transitions (Figure 5I).

## Discussion

The substantial redundancy within the TPX family leads to weak single loss-of-function phenotypes, and may obscure underlying functional differences (Causier et al., 2012c; Long et al., 2006; Plant et al., 2021). Using higher order mutants in combination with immunity challenges and auxin-regulated developmental readouts, we discovered subclade-specific roles, and found cross-regulation within the family, namely direct repression of *TPR2* and *TPR3* expression by *TPL* and *TPR1*. Higher order mutants impacted phenotypes across the plant, including: number of leaves, time to flowering, response to pathogen infection, and root and floral organ development. Transcriptome analysis reflected the trends observed in organism-level phenotypes and showed that stress responses exaggerated functional divergence within the family. Synthetic repression assays showed that variation in the EAR-binding domain results in weaker activity by *TPR2* and *TPR3* compared to the other family members. Expression of EAR-bearing pathogen effectors like RxL21 created sensitized conditions in our synthetic repression assay that further distinguished client recruitment activity among family members, while also successfully lowering the threshold for auxin response in plants. Together, our results support a model where TPX function is well-buffered under normal conditions but becomes sensitive to reductions in dosage during cell state transitions like infection or organogenesis. Additionally, these sensitivity points highlight how the graded affinity between TPX proteins and their EAR-containing client proteins may have been shaped by evolving affinities reflective of the TPX evolutionary tree.

The functional split between *TPL/TPR1/TPR4* and *TPR2/TPR3* is most sharply defined in the context of plant immunity. Our transcriptomic analysis following *B. cinerea* inculcation revealed that *TPR2* and *TPR3* contributed to regulation of gene sets distinct from those controlled by *TPL/TPR1/TPR4*. Griebel et al. (2023) demonstrated that *TPR1* together with *TPL* and *TPR4*, supports initial immune activation but subsequently mitigates adverse effects of prolonged defense signaling on host physiology and growth. Multiple oomycete, fungal, bacterial, and insect effectors target TPX proteins through diverse mechanisms (Bindics et al., 2022; Darino et al., 2021; Downing et al., 2025; Harvey et al., 2020; Navarrete et al., 2022b, 2021), and the *Arabidopsis*–Hpa interactome reveals extensive effector engagement with host corepressor networks (Mukhtar et al., 2011; Wessling et al., 2014). The convergence of effectors from across the tree of life on the TPX interface underscores the need for full TPX dosage when cells undergo large-scale transcriptional reprogramming. The observation that competition is strongest in the H8-23 (TPR2/TPR3) background directly connects corepressor sequence variation to effector susceptibility and is consistent with the infection data in mutant plants, particularly in light of the many candidate *B. cinerea* effector proteins that contain EAR sequences.

One of the first identified and best studied clients of the TPX family are the Aux/IAA proteins. Recently, some Aux/IAAs were shown to exhibit bimodal behavior, where depending on the auxin concentration in the cell, they could switch between activator and repressor modes (Cho et al., 2024). A single amino acid swap in the EAR motif (from LRLGL to LCLGL) is sufficient to convert a bimodal-type Aux/IAA to consistent repression and is correlated with relative affinity for TPL. Our findings complement this model by revealing an analogous source of variation on the corepressor side. The Helix 8 sequence of TPR2 and TPR3 reduces client recruitment efficiency without eliminating repression capacity, effectively lowering the affinity of the corepressor for its EAR-bearing client. Within a bimodal framework, the same Aux/IAA protein could produce different transcriptional outcomes depending on the complement of TPX proteins available: stronger paralogs would stabilize the repressed state across a wider range of Aux/IAA concentrations, whereas TPR2 and TPR3 would permit greater variability or partial de-repression. This framework provides a mechanistic hypothesis for why family members appear redundant under some conditions but are unable to compensate fully for one another during some critical challenges, like development of root hairs or exposure to *B. cinerea*. Specifically, our results suggest that the absolute level of TPX function needed to maintain repression increases to a different extent during different cell state transitions and/or that different affinities between each TPX family member and potential clients impacts which repression complexes can be maintained when TPX levels are limiting.

Current tools for engineered repression rely on recruitment of endogenous TPX complexes through synthetic EAR domains (SRDX; (Hiratsu et al., 2003) or direct fusion of the TPL N-terminal domain to transcription factors (HACR; (Khakhar et al., 2018). Helix 8 as a tunable determinant of recruitment efficiency suggests that engineered variants could provide a graded set of repression strengths without altering the repressive machinery itself. Combined with the bimodal properties of different EAR motif sequences (Cho et al., 2024), this creates a two-dimensional design space for specifying transcriptional outputs; one axis defined by the client EAR motif and the other by the corepressor Helix 8 sequence (Downing et al., 2025; Leydon et al., 2020; Markel et al., 2024). The cross-regulation of expression within the family also hints at a more universal strategy for balancing robustness and tunability in multigene corepressor families, as there are clear parallels to Groucho/TLE corepressors in animals (Agarwal et al., 2015; Jennings and Ish-Horowicz, 2008).

## Figure Supplements

**Figure 1 - figure supplement 1.**
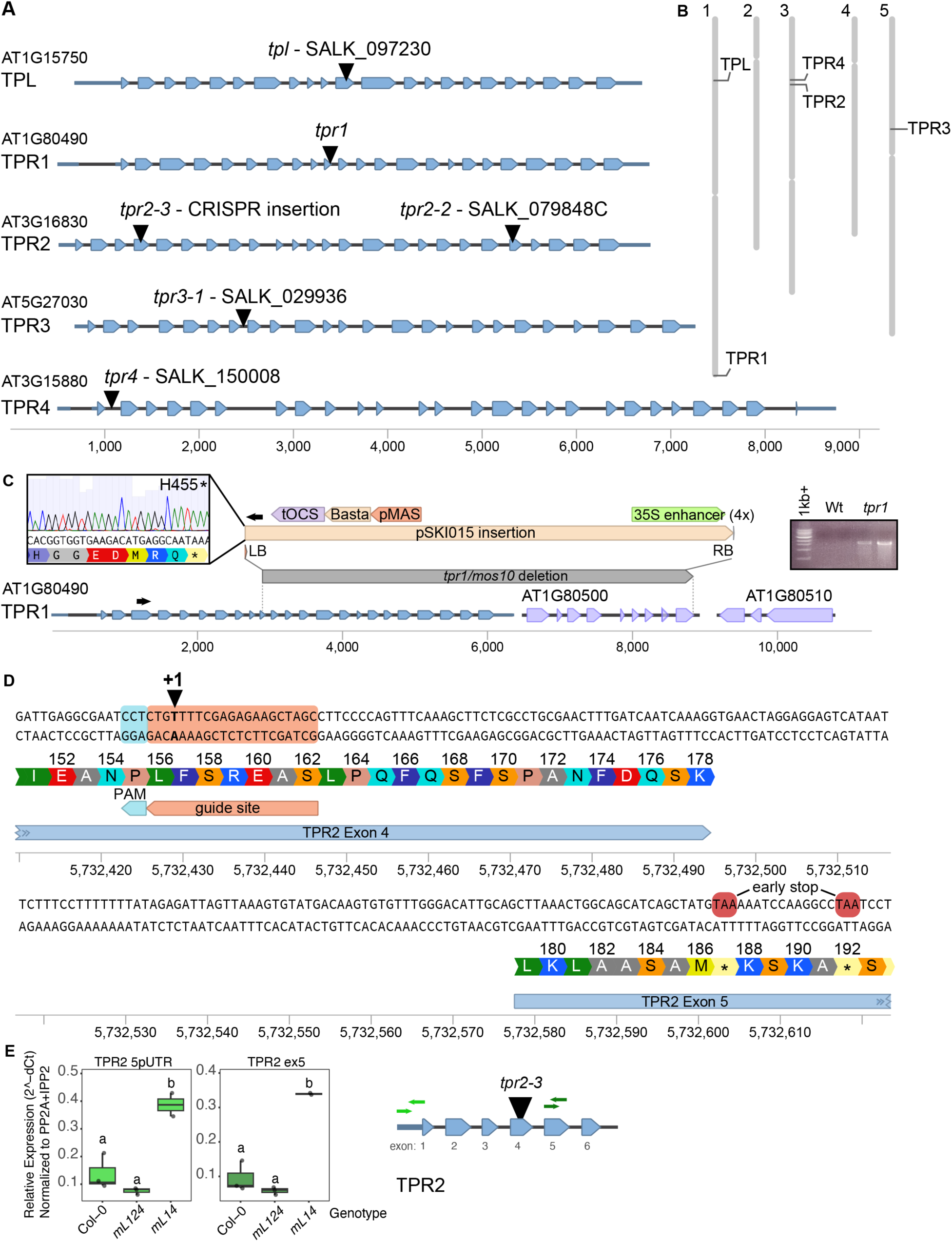
TPX gene models, mutant alleles, and CRISPR design. **(A)** Gene models for *TPL* (SALK_097230), *TPR1* (T-DNA induced deletion), *TPR2* (*tpr2-3* CRISPR insertion; *tpr2-2* SALK_079848C), *TPR3* (SALK_029936), *TPR4* (SALK_150008). Scale in bp. **(B)** Chromosomal positions of all five genes. **(C)** Detailed sequence of the *tpr1-1*/*mos10* allele showing pSKI015 insertion, orientation, left and right borders. Electrophoresis gel of targeted amplification of the *tpr1* allele, with associated Sanger sequencing shown in the inset: the allele induces an early stop at H455. **(D)** Detailed sequence of the *tpr2-3* CRISPR allele showing guide site at the Exon 4/5 junction, amino acid positions 152–192, and premature stop codons introduced within the CRA domain. **(E)** qPCR with four primer sets, upstream and downstream of the CRISPR insertion site, confirms reduction of *TPR2* expression in *tpr2-3*.

**Figure 2 - figure supplement 1.**
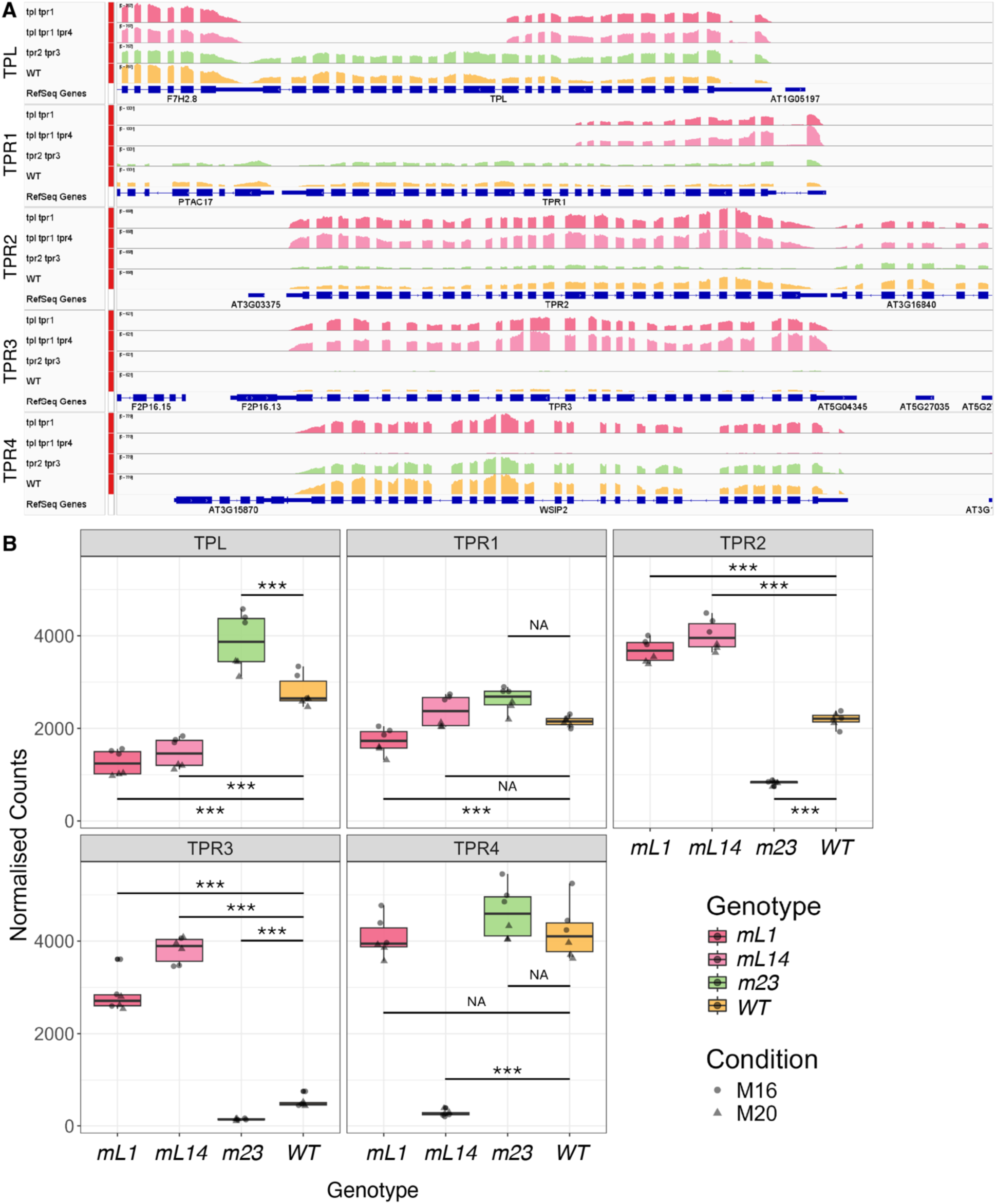
Molecular characterization of the impact of TPX mutants on family member expression by RNA-Seq. **(A)** Sequence tracks for each of the 5 TPX family members (TPL, TPR1-4) analyzed in multiple mutant backgrounds. These results confirm an absence of full-length mRNA for each of the T-DNA based mutants defined in Figure 1 - figure supplement 1. The *tpr1* allele contains a deletion (see Figure 1 – figure supplement 1), and this results in no detectable RNA downstream of the deletion site detectable by RNA-seq. RNA levels from the residual locus in *tpr1* mutants are unchanged in aggregate compared to wild type, which suggests upregulation of the transcript in the absence of full length TPR1 protein. **(B)** Quantification of total normalized counts over each gene body plotted as boxplots. *** indicates a significant difference between genotypes (p<0.01). NA – no significant difference. These results support the cross-family regulation observed by RT-qPCR in Figure 1.

**Figure 2 - figure supplement 2.**
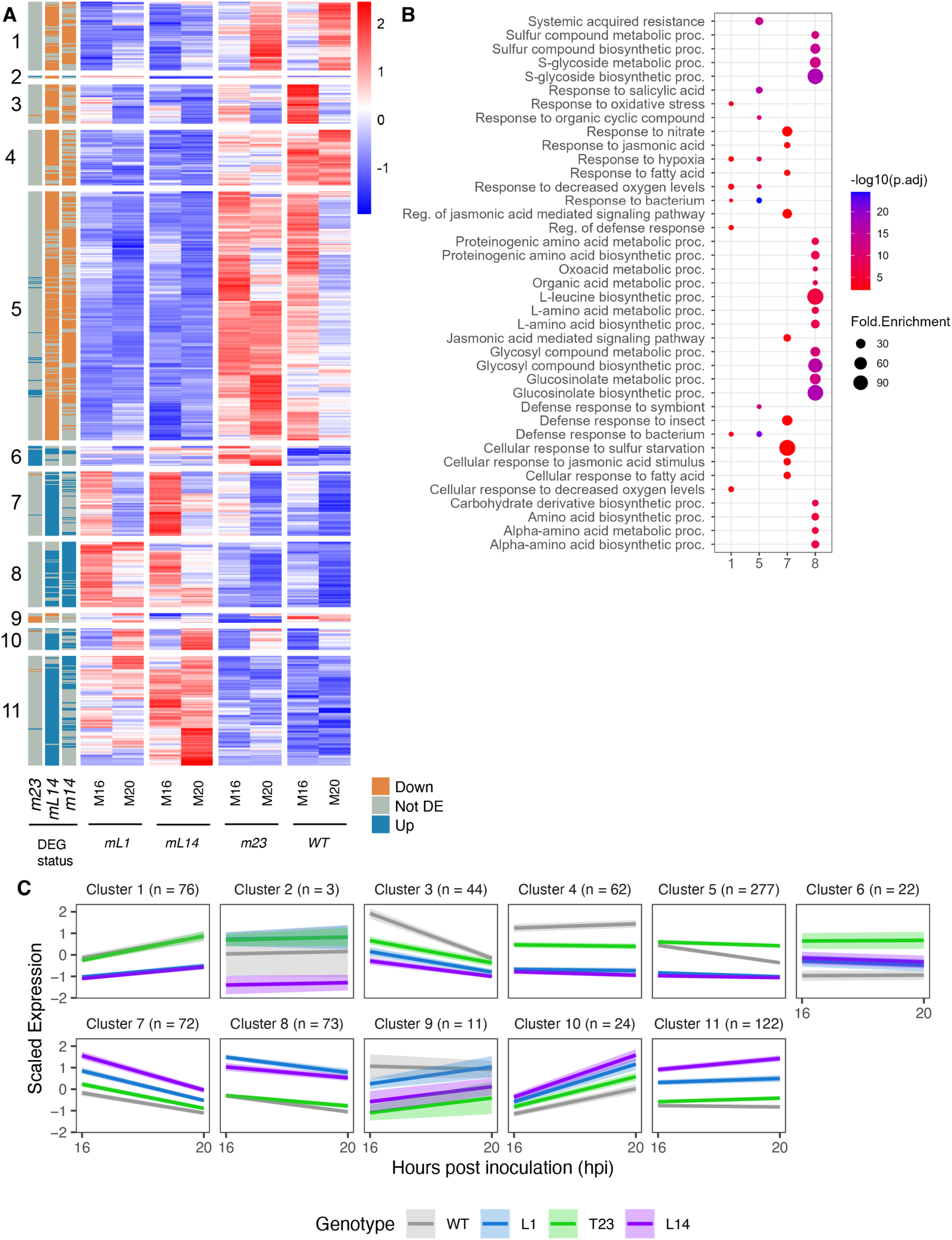
TPX family multiple mutants demonstrate a semi-overlapping impact on the leaf transcriptome. **(A)** Heatmap showing row-scaled (z-score) normalized expression of differentially expressed genes (DEGs) identified in mL1, mL14, and m23 relative to wild type under mock conditions (padj < 0.01, |log2FC| ≥ 0.5). Columns represent mean normalized counts ordered by genotype (WT, mL1, m23, mL14) at 16 and 20 hours post inoculation (hpi). Row annotations indicate whether a gene is significantly differentially expressed in *mL1, mL14,* and *m23* compared to WT (Downregulated is blue; Upregulated is orange, not differentially expressed is grey). **(B)** Gene Ontology Biological Process enrichment of DEGs from each transcriptional module identified in (A). **(C)** Mean scaled expression profiles (z-score) of genes within each of the 11 transcriptional modules defined by hierarchical clustering of DEGs in *mL1, mL14, and m23* relative to WT (see A), across 16 and 20 hpi under mock conditions. Lines are colored by genotype (WT, gray; mL1, blue; m23, green; mL14, purple) and shaded ribbons indicate 95% confidence intervals.

**Figure 2 - figure supplement 3.**
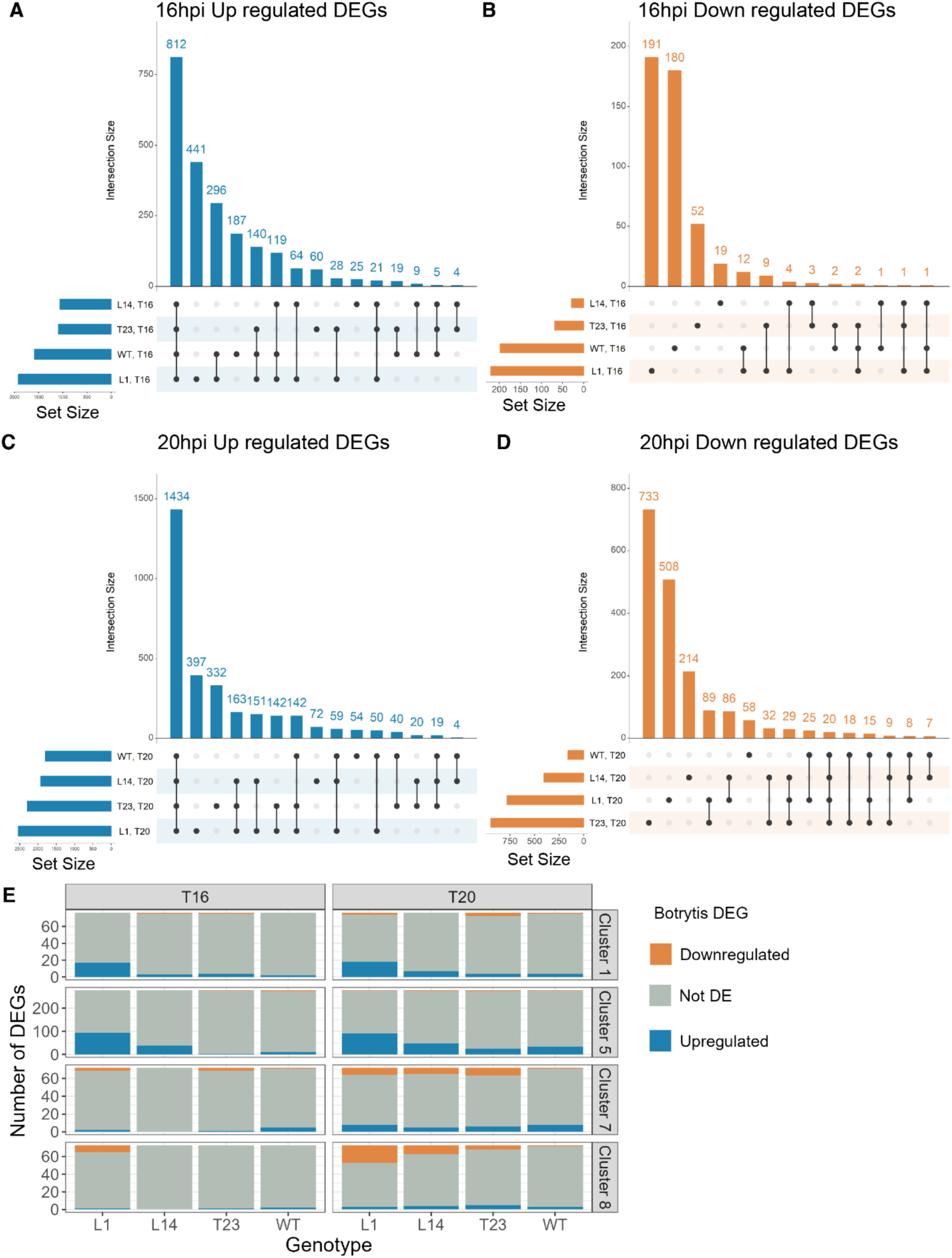
UpSet plots showing overlap of differentially expressed genes in response to *Botrytis cinerea* infection across genotypes. UpSet plots showing the overlap of upregulated **(A, C)** and downregulated **(B, D)** differentially expressed genes (DEGs) identified in response to *B. cinerea* infection relative to mock (padj < 0.01, |log2FC| ≥ 0.5) in wild type (WT), *tpl tpr1* (*mL1*), *tpl tpr1 tpr4* (*mL14*), and *tpr2 tpr3* (*m23*) at 16 hpi **(A, B)** and 20 hpi **(C, D)**. Bar height indicates the number of DEGs unique to or shared between each genotype combination. Set size bars indicate the total number of DEGs per genotype. **(E)** Bar graph showing the genes in four transcriptional modules of DEGs in *tpl tpr1* (*mL1*), *tpl tpr1 tpr4* (*mL14*), and *tpr2 tpr3* (*m23*) mutants compared to wild type (WT) from the mock data (Figure 2 – figure supplement 2) and whether the same genes are differentially expressed in each of the genotypes in response to *B. cinerea* expression at 16 and 20 hours post infection. Orange represents genes that are downregulated upon *B. cinerea* infection, Blue represents genes that are upregulated in response to *B. cinerea* infection, and Grey represents genes that are not significantly differentially expressed in each genotype in response to *B. cinerea* infection.

**Figure 2 Figure Supplement 4.**
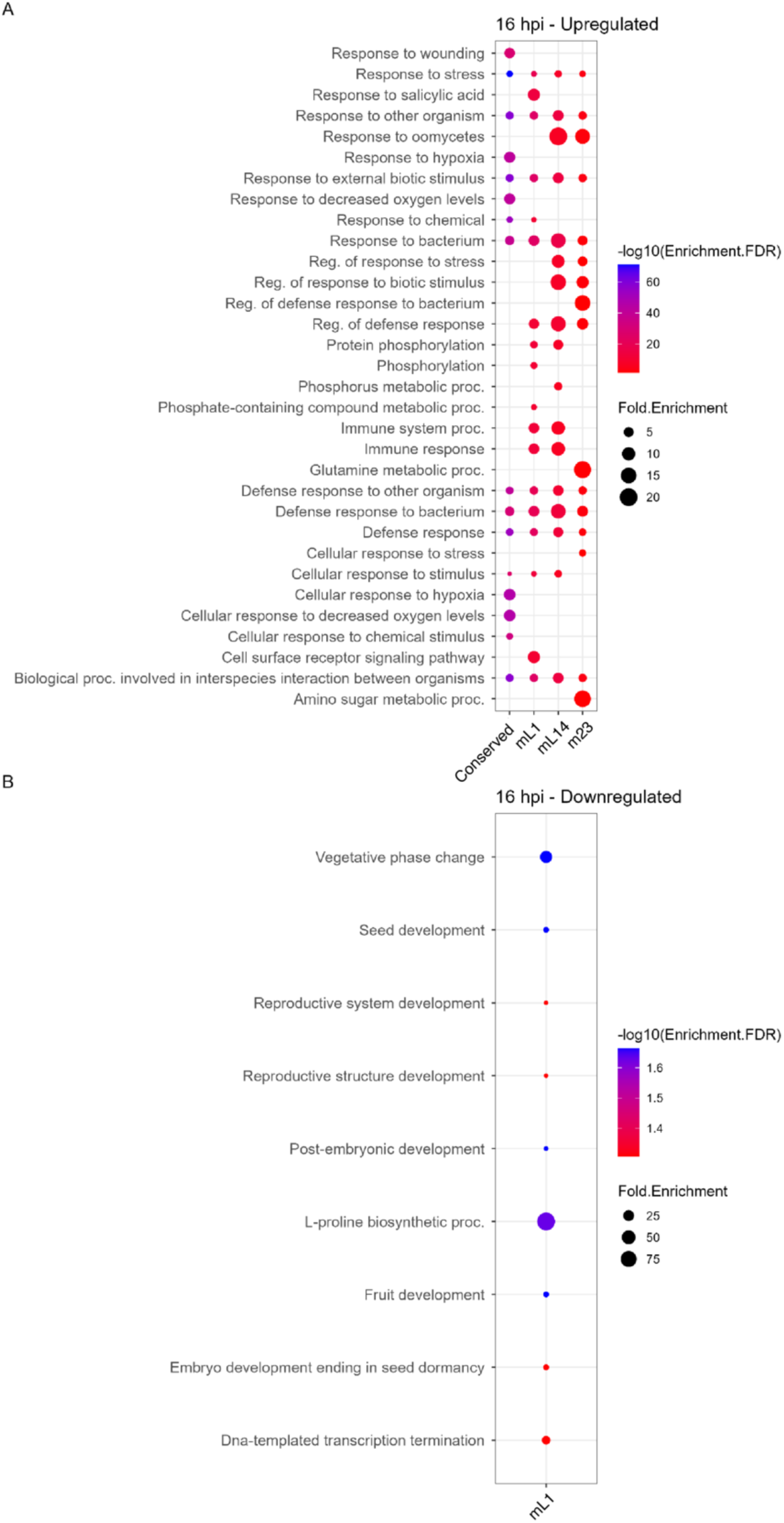
GO term analysis for conserved and mutant-specific DEGs after *B. cinerea* infection at 16 hpi. **(A)** GO term analysis for conserved (n = 1400), *tpl tpr1* specific (*mL1*, n = 554), *tpl tpr1 tpr4* specific (*mL14*, n = 114), and *tpr2 tpr3* specific (*m23*, n = 113) upregulated DEGs at 16 hpi. **(B)** GO term analysis for conserved (n = 18), *tpl tpr1*specific (*mL1*, n = 205), *tpl tpr1 tpr4* specific (*mL14*, n = 27), and *tpr2 tpr3* specific (*m23*, n = 65) downregulated DEGs at 16 hpi. Conserved genes were defined as being expressed in WT and at least one of the mutants, in the same direction at the same timepoint; mutant-specific genes were defined as being differentially expressed in the mutant but not in WT, at the same timepoint.

**Figure 2 Figure Supplement 5.**
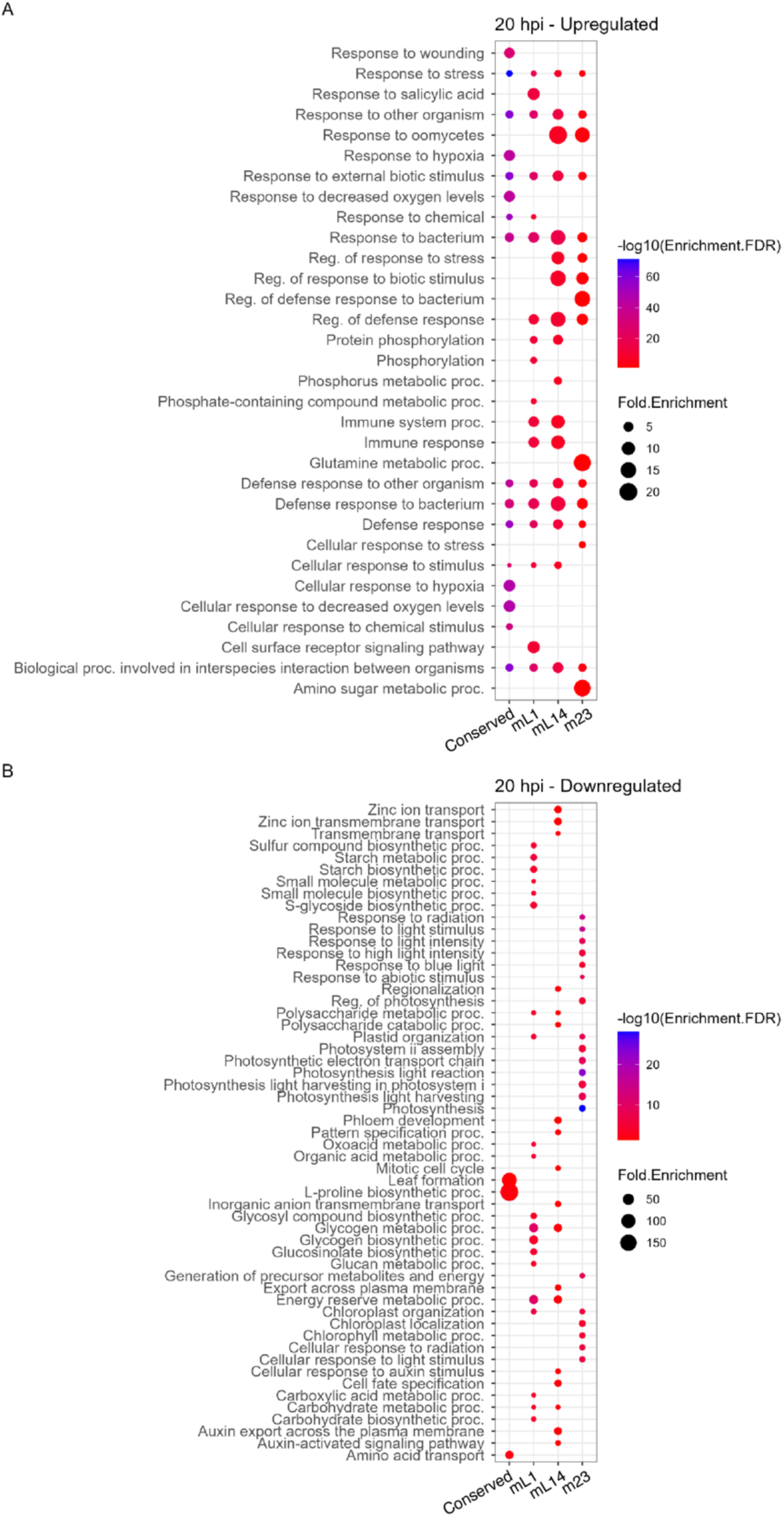
GO term analysis for conserved and mutant-specific DEGs after *B. cinerea* infection at 20 hpi. **(A**) GO term analysis for conserved (n = 1748), *tpl tpr1* specific (*mL1*, n = 853), *tpl tpr1 tpr4* specific (*mL14*, n = 406), and *tpr2 tpr3* specific (*m23*, n = 657) upregulated DEGs at 20 hpi. **(B)** GO term analysis for conserved (n = 102), *tpl tpr1*specific (*mL1*, n = 712), *tpl tpr1 tpr4* specific (*mL14*, n = 361), and *tpr2 tpr3* specific (*m23*, n = 883) downregulated DEGs at 20 hpi. Conserved genes were defined as being expressed in WT and at least one of the mutants, in the same direction at the same timepoint; mutant-specific genes were defined as being differentially expressed in the mutant but not in WT, at the same timepoint.

**Figure 3 - figure supplement 1.**
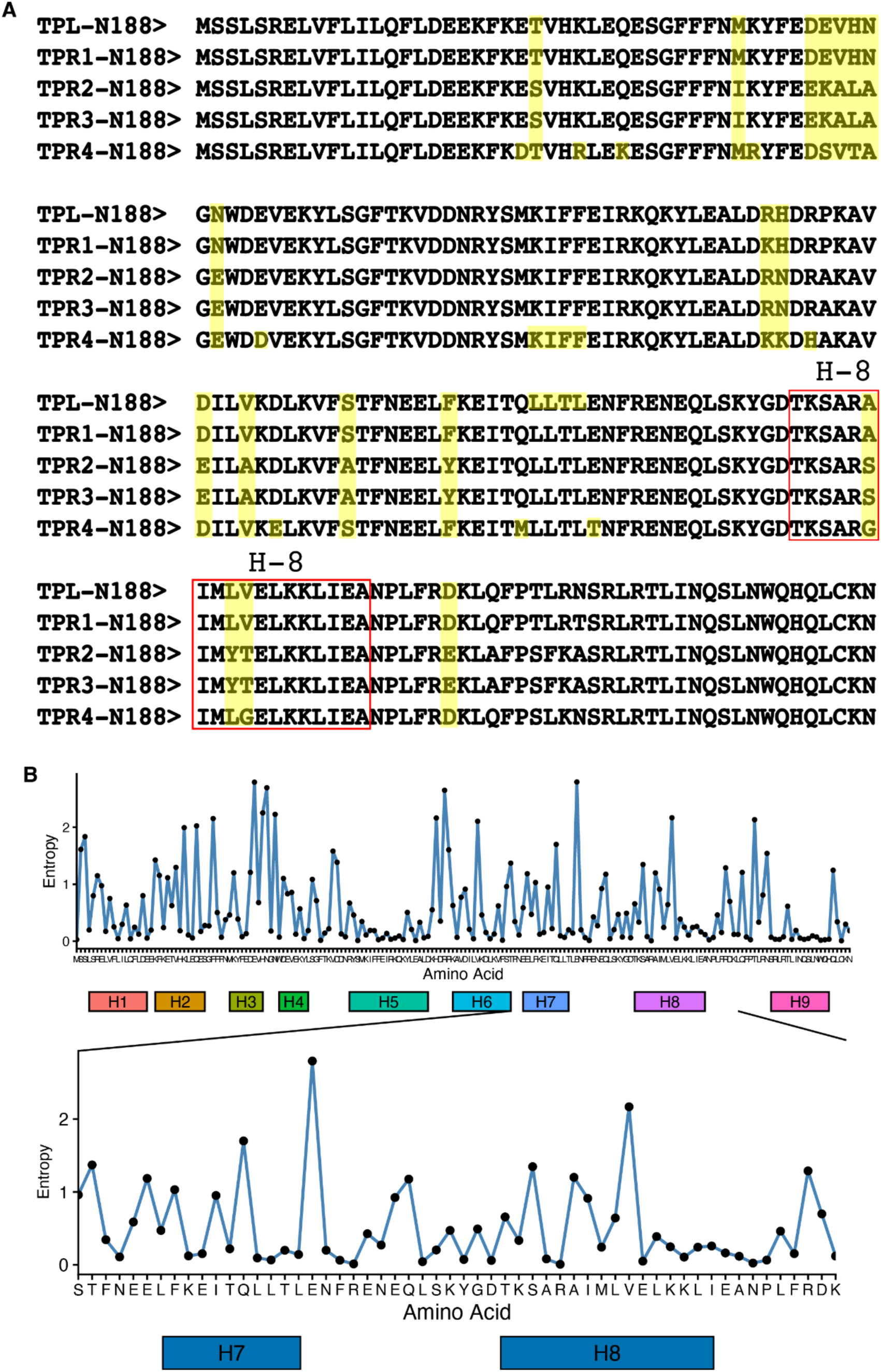
Amino acid sequence alignment and variation in TPX family proteins. **(A)** Amino acid alignment of TPL and TPR1-4. Alignment performed in MEGA, variant residues are highlighted in yellow. Helix 8 region is denoted with a red box. **(B)** Per-residue sequence entropy along the protein, computed with PoET. Sequence divergence was quantified using PoET (Protein Evolutionary Transformer), an autoregressive, retrieval-augmented protein language model from OpenProtein.AI that models whole protein families as sequences-of-sequences without requiring a multiple sequence alignment. Conditioned on a homologous prompt set drawn from the target family, PoET returns a per-position log-likelihood over the 20 amino acids, which we converted to relative frequencies and summarized as Shannon entropy at each residue. Entropy (y-axis) is plotted against residue position (x-axis) across the full reference sequence. Peaks indicate positions tolerant of substitution and representative of diverged/variable sites (“mutational hotspots”), whereas troughs mark evolutionarily constrained, conserved positions. Analysis was performed on openprotein.ai servers (Truong and Bepler, 2023)

**Figure 3 - figure supplement 2.**
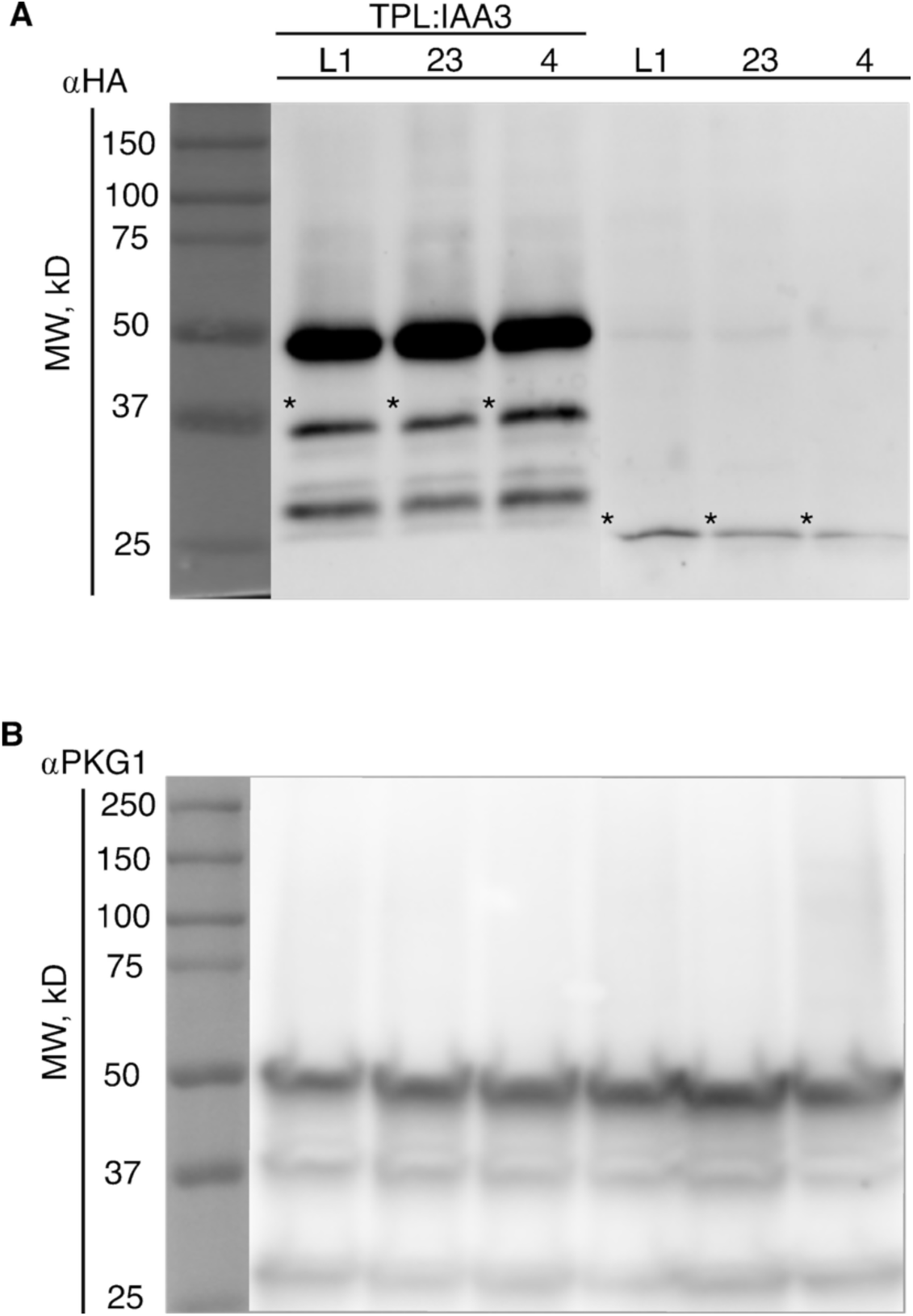
TPL H8 EAR binding pocket cytometry protein expression. **(A)** ⍺HA Western blots of TPL H8 EAR subtype binding pocket cytometry stains blotting for TPL expression. Unfused or Free variants of L1, 23, and 4 primary band noted with asterisks (MW, KD = ∼24), Fused IAA3 variants primary band noted with asterisks (MW, KD = ∼ 36). **(B)** ⍺PkG1Western blots of TPL H8 EAR subtype binding pocket cytometry stains blotting for control yeast protein expression (MW, KD = ∼45).

**Figure 3 - figure supplement 3.**
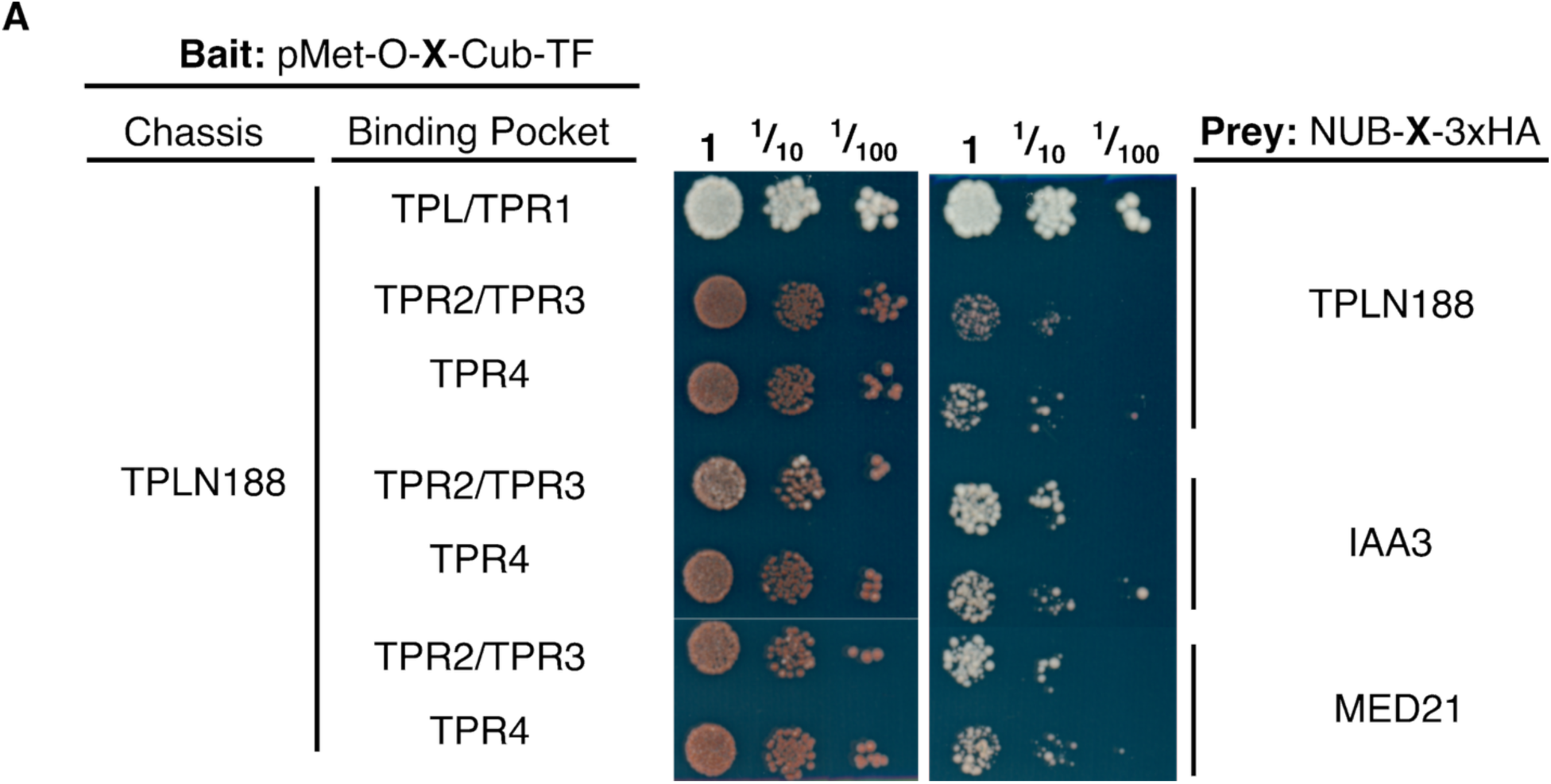
TPX H8 variation effects on corepressor client binding. **(A)** Growth control test (right) and interaction test (left) CytoSUS plates of H8 TPL variants (23 and 4) to test for physical interaction with known TPL interactors TPLN188, IAA3, and MED21

**Figure 4 - figure supplement 1.**
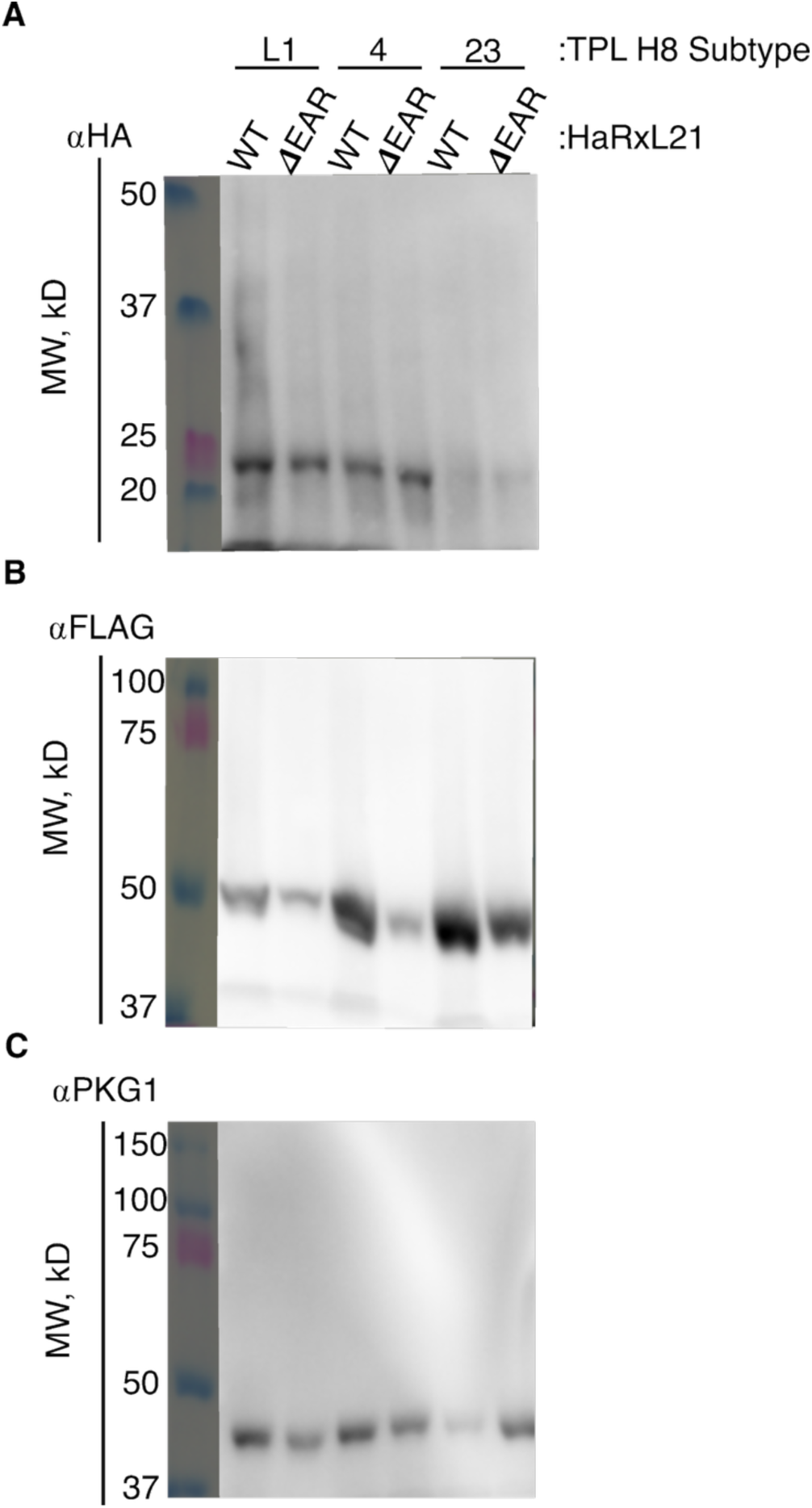
TPX protein expression in RxL21 competition assays. **(A)** ⍺HA Western blots of TPL H8 EAR subtype binding pocket with RxL21 (WT and *Δ*EAR) protein extracted from cytometry stains blotting for TPL expression. Subtypes L1, 23, and 4 primary band (MW, KD = ∼24). **(B)** ⍺FLAG Western blots of TPL H8 EAR subtype binding pocket with RxL21 (WT and *Δ*EAR) cytometry stains blotting for RxL21 (WT or *Δ*EAR) protein expression. RxL21 Wt and *Δ*EAR primary band (MW, KD = ∼49). **(C)** ⍺PkG1Western blots of TPL H8 EAR subtype binding pocket with RxL21 (WT and *Δ*EAR) cytometry stains blotting for control yeast protein expression (MW, KD = ∼45).

**Figure 4 - figure supplement 2.**
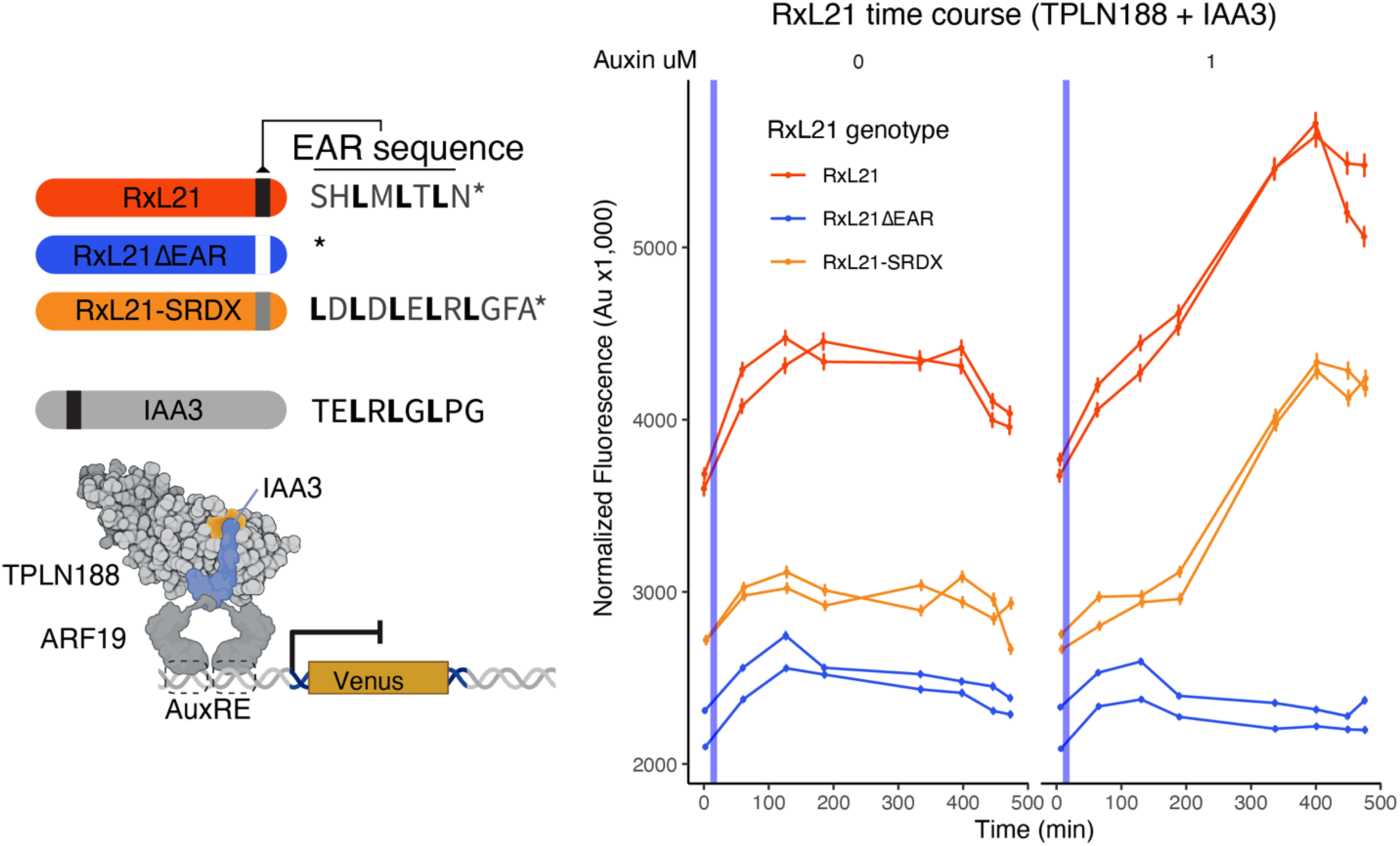
Auxin effects on RxL21 competed TPX repression. Time course flow cytometry of previously tested RxL21 competition strains with TPLN188 (Figure 5B) with the addition of Auxin (vertical blue bar) to further sensitize the system to RxL21 disruption by promoting the degradation of IAA3. Wild type RxL21 (Red), RxL21 with deletion of the C-terminal EAR motif (blue), and replacement of the native EAR sequence with the SRDX EAR sequence (orange) strains were treated with either no auxin (mock, left panel), or 1µM IAA (right panel), and cytometry was performed over the following 500 minutes. Every point represents the average fluorescence of 5–10,000 individually measured yeast cells (a.u.: arbitrary units). Auxin (IAA-10 µM) was added at the indicated time (gray bar, +Aux). Error bars represent standard error.

**Figure 4 - figure supplement 3.**
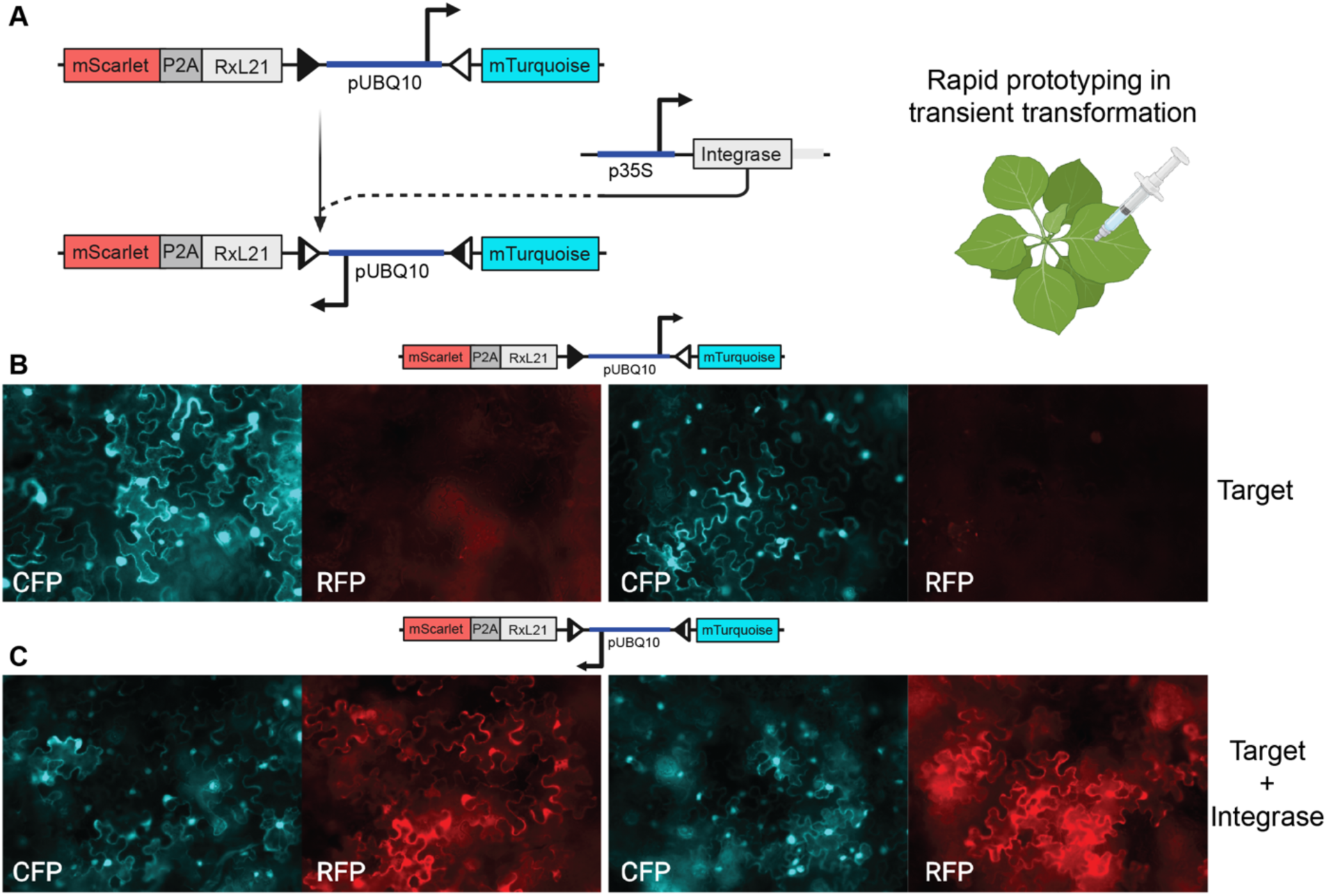
Rapid prototyping of iInducer constructs in transient transfections of *Nicotiana benthamiana* at 2 days after injection. **(A)** Design of the integrase target. The target is composed of two PhiC31 integrase sites (triangles) surrounding a constitutive promoter (pUBQ10), the fluorescent reporter mTurquoise and the RxL21 CDS linked to a P2A cleavage sequence and a fluorescent reporter (mScarlet). In the absence of integrase mTurquoise is expressed. In the presence of integrase, the integrase (PhiC31) mediates inversion of the DNA between the integrase sites, inverting the promoter and leading to RxL21 and mScarlet expression. The expression of the integrase is mediated by the selected promoter selected, here the viral promoter p32S. **(B)** On the left side is the wild-type RxL21-P2A-mScarlet target that switches from mTurquoise to RxL21-P2A-mScarlet alone (top) and **(C)** with a *p35S:PhiC31* construct (bottom). **(B)** On the right side shows the RxL21ΔEAR-P2A-mScarlet target alone (top) and (**C)** RxL21ΔEAR-P2A-mScarlet target with a *p35S:PhiC31* construct (bottom). Microscopy images were taken on a 20x objective to allow a wide view of switching efficiency.

**Figure 5 - figure supplement 1.**
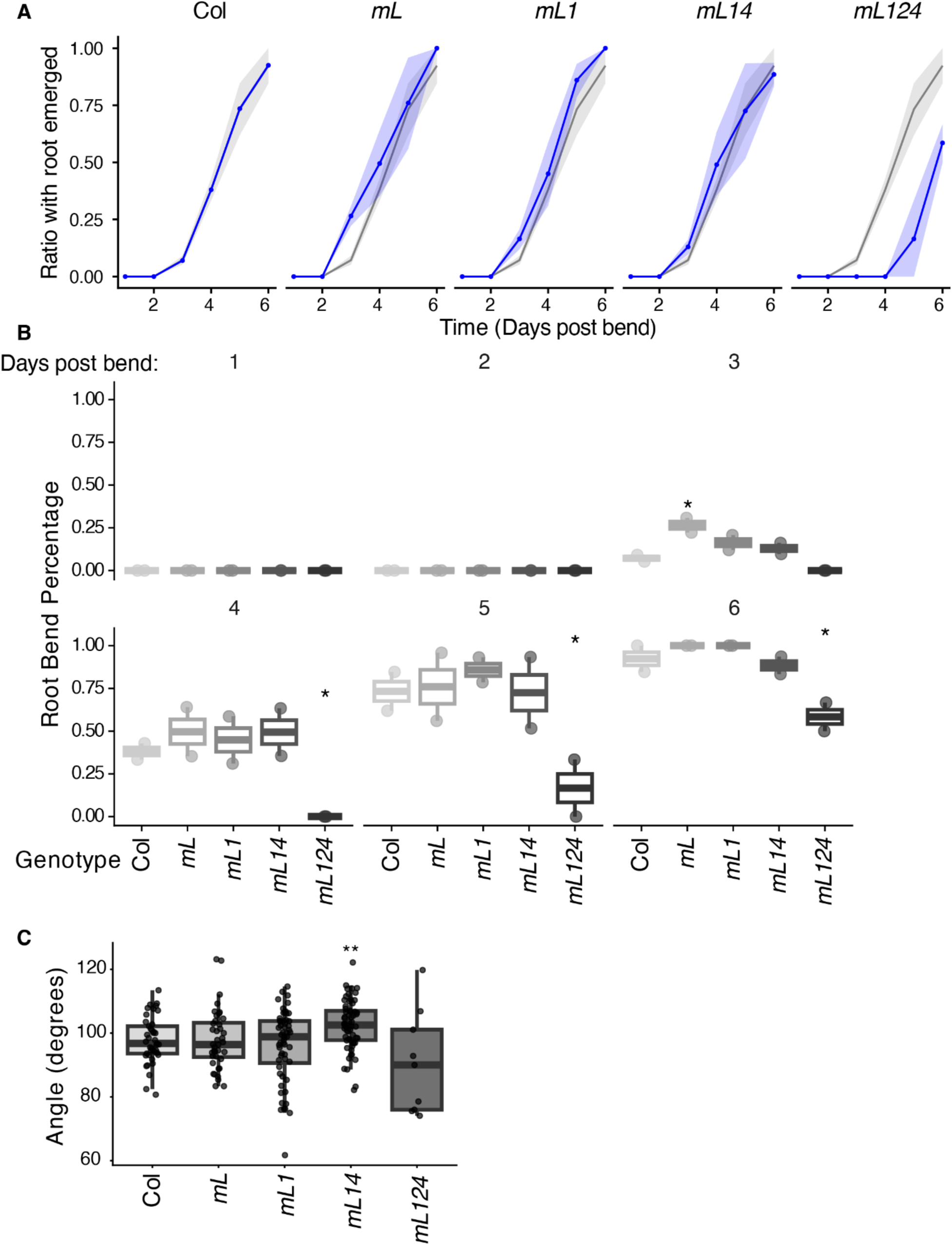
Root phenotypes in TPX multiple mutants. (A-B) Time course of lateral root emergence at the bend site. Mean proportion of seedlings for Col, *tpr1 (m1)*, *tpl tpr1 (mL1)*, *tpl tpr1 tpr4 (mL14)*, and *tpl tpr1 tpr2 tpr4 (mL124)* genotypes across 1–6 days post-bend (dpb), with an emerged lateral root at the bend site plotted over 1–6 days post-bend (dpb) for each genotype. **(A).** Each panel shows an individual genotype (blue line and shaded ribbon, indicating the range across 2 plates) overlaid on the Wild-type (Col-0) reference (grey line and shaded ribbon). Seedlings were rotated 90° to induce gravitropic bending and scored daily. n = 2 plates per genotype per timepoint. **(B)** Box plot - boxes represent the interquartile range, with the median shown as a horizontal line; individual plate values are overlaid as points. Significance relative to Col was assessed at each day using Fisher’s exact test on pooled counts across plates, with Benjamini-Hochberg correction for multiple comparisons. * p < 0.05. n = 2 plates per genotype per timepoint. **(C)** Root bend angles across genotypes. Angles of the primary root at the bend site measured from individual seedlings at 2 days post-bend for Col, *tpr1 (m1)*, *tpl tpr1 (mL1)*, *tpl tpr1 tpr4 (mL14)*, and *tpl tpr1 tpr2 tpr4 (mL124)* genotypes. Boxes represent the interquartile range with the median shown as a horizontal line; individual root measurements are overlaid as points. Angles were measured from TIFF images using ImageJ. Significance relative to Col was assessed using a Wilcoxon rank-sum test with Benjamini-Hochberg correction for multiple comparisons. ** p < 0.01. n = Col: 41, *tpr1*: 40, *tpl tpr1*: 56, *tpl tpr1 tpr4*: 62, *tpl tpr1 tpr4 tpr2*: 9 roots across 2 plates.

**Figure 5 - figure supplement 2.**
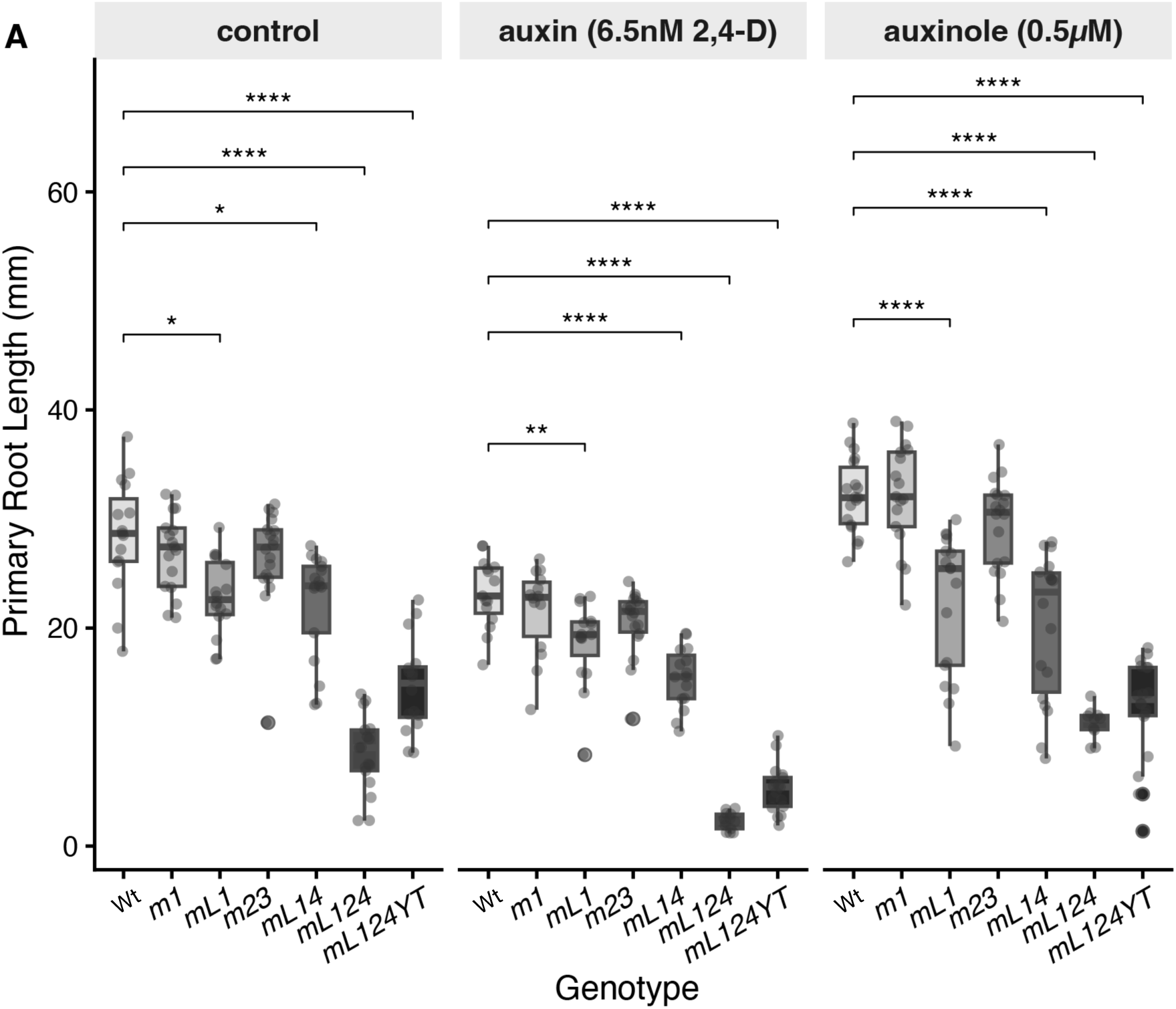
Primary root length of *Arabidopsis thaliana* TPX mutants in response to auxin and auxinole. 9A-B) Primary root length of wild-type (Col-0) and TPX mutants Col-0 (WT), *tpr1* (*m1*), *tpl tpr1* (*mL1), tpr2 tpr3* (*m23*), *tpl tpr1 tpr4* (*mL14*), *tpl tpr1 tpr2 tpr4* (*mL124*) and *tpl tpr1 tpr2 tpr4 YFP:TPL* (*mL124YL*) seedlings grown vertically on ½ LS medium supplemented with 6.5 nM 2,4-D (auxin) or 0.5 µM auxinole for 4 days. Seedlings were photographed and root length measured using Fiji. Boxplots show the median (center line), interquartile range (box), and 1.5× IQR (whiskers); individual measurements are overlaid as points. Statistical comparisons between each genotype and Col-0 within each treatment were performed using a two-tailed Student’s t-test with Bonferroni correction. Significance is indicated as: * p ≤ 0.05, ** p ≤ 0.01, *** p ≤ 0.001, **** p ≤ 0.0001; ns, not significant. n = 18 seedlings per genotype per treatment pooled from 3 independent experiments.

**Figure 5 - figure supplement 3.**
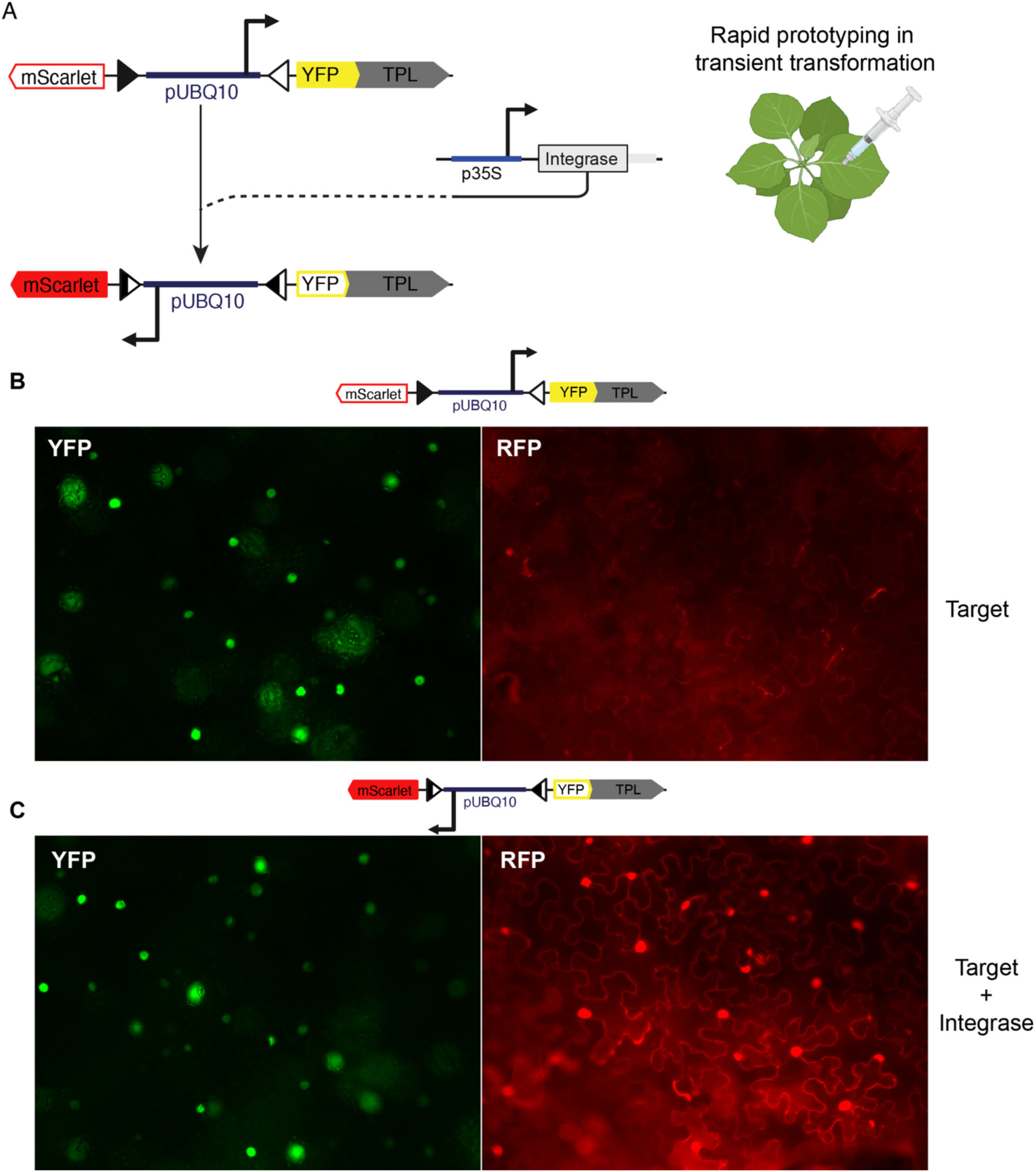
Rapid prototyping of iEraser constructs in transient transfections of *Nicotiana benthamiana* at 2 days after injection. **(A)** Design of the integrase target. The target is composed of two PhiC31 integrase sites (triangles) surrounding a constitutive promoter (pUBQ10), the YFP-tagged TPL full length CDS and a fluorescent reporter (mScarlet). In the absence of integrase YFP-TPL is expressed. In the presence of integrase, the integrase (PhiC31) mediates inversion of the DNA between the integrase sites, inverting the promoter and leading to mScarlet expression. The expression of the integrase is mediated by the selected promoter selected, here the viral promoter p32S. **(B)** iEraser target expression in the absence of integrase. **(C)** iEraser target expression in the presence of a *p35S:PhiC31* construct. On the left side. On the left side is the YFP channel for YFP-TPL, and mScarlet is on the right side. Microscopy images were taken on a 20x objective to allow a wide view of switching efficiency.

**Figure 5 - figure supplement 4.**
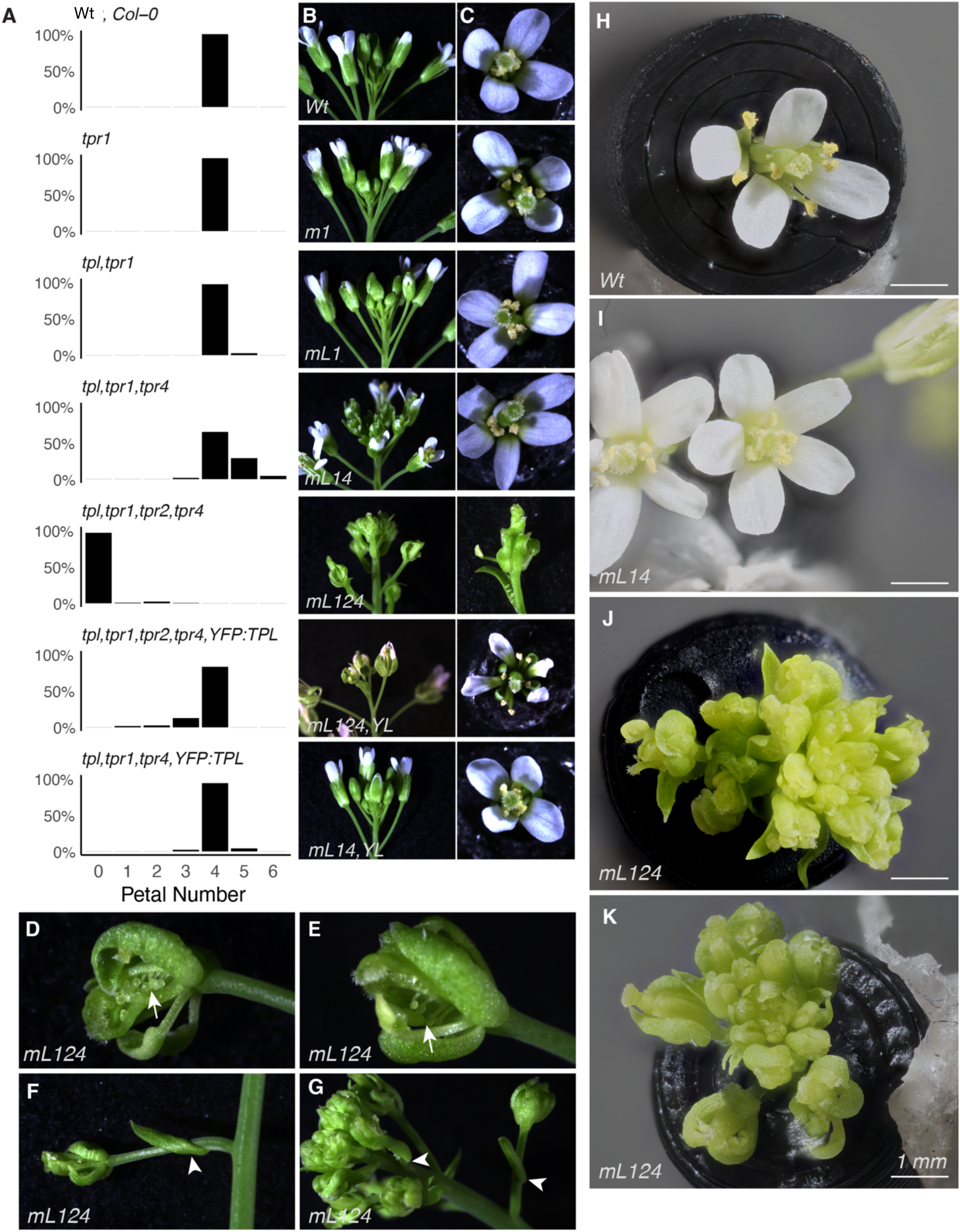
Flower phenotypes of TPX mutants. **(A)** Flower petal quantification of 150 flowers for each genotype data is presented as the ratio of the total. Wild-type (Col-0) and TPX mutants Col-0 (WT), *tpr1* (*m1*), *tpl tpr1* (*mL1), tpr2 tpr3* (*m23*), *tpl tpr1 tpr4* (*mL14*), *tpl tpr1 tpr2 tpr4* (*mL124*) and *tpl tpr1 tpr2 tpr4 YFP:TPL* (*mL124YL*) **(B)** Column is inflorescence **(C)** Column is individual flowers from above **(D-E)** Highlighting ectopic ovule formation (arrow). **(F-G)** Highlight novel induction of bracts (arrow). **(H-K)** MacroPod PRO images for high resolution focus stacked observation of floral morphology in Wt, *mL14*, and *mL124*.

**Figure 5 - figure supplement 5.**
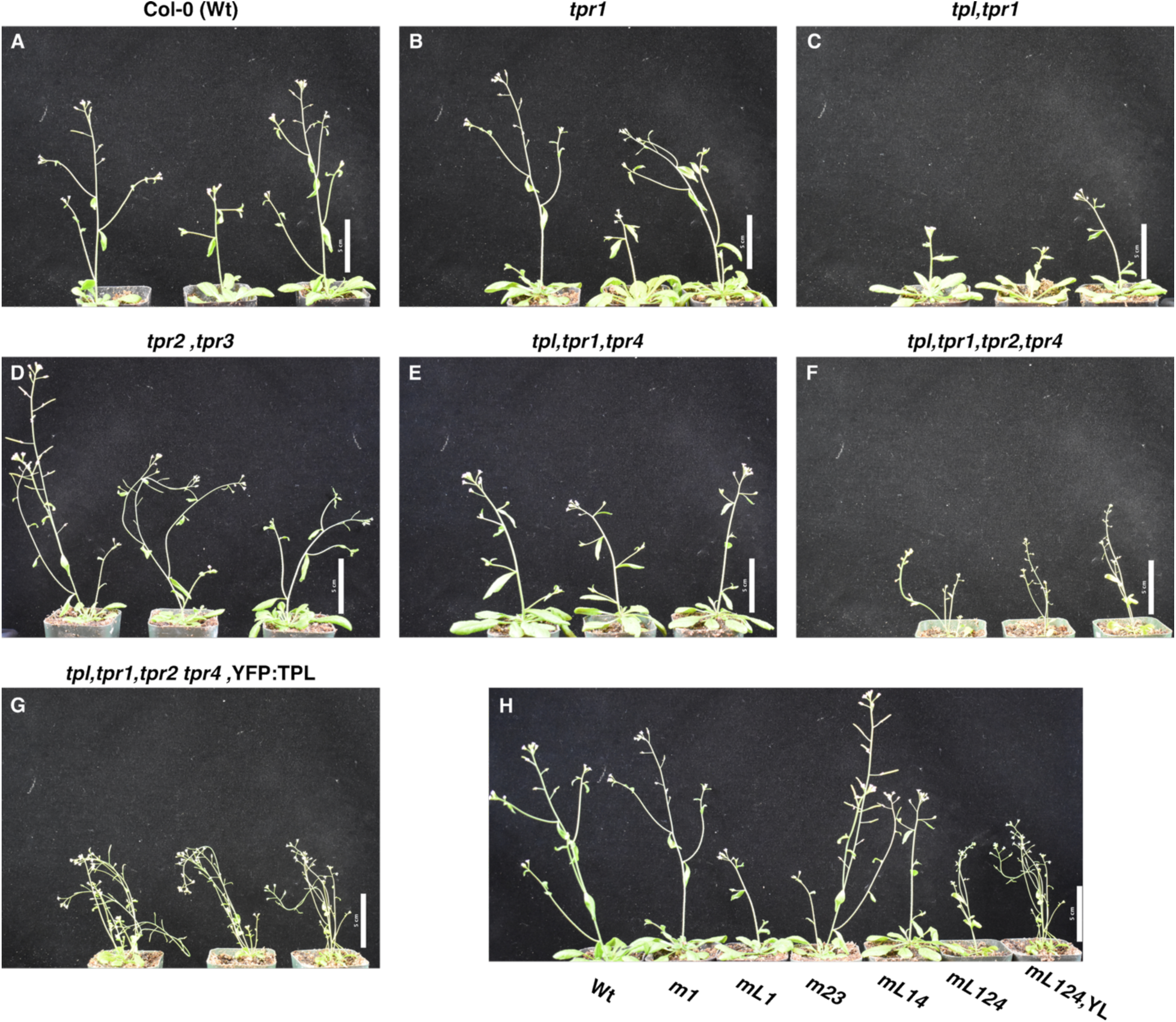
Whole plant phenotypes of TPX mutants. **(A)** Three whole plant representatives of wild-type Col-0 (Wt) genotype with a 5 cm. scale bar. **(B)** Three whole plant representatives of *tpr1* (*m1)* genotype with a 5 cm. scale bar. **(C)** Three whole plant representatives of *tpl tpr1* (*mL1)* genotype with a 5 cm. scale bar. **(D)** Three whole plant representatives of *tpr2 tpr3* (*m23*) genotype with a 5 cm. scale bar. **(E)** Three whole plant representatives of *tpl tpr1 tpr4* (*mL14*) genotype with a 5 cm. scale bar. **(F)** Three whole plant representatives of *tpl tpr1 tpr2 tpr4* (*mL124*) genotype with a 5 cm. scale bar. **(G)** Three whole plant representatives of *tpl tpr1 tpr2 tpr4 YFP:TPL* (*mL124,YL*) genotype with a 5 cm. scale bar. **(H)** Full plant images of representative member of each genotype Wt and TPX mutants: *m1*, *mL1, m23*, *mL14*, *mL124*, and *mL124,YL* with a 5 cm. scale bar.

## Materials and Methods

### Key Resources Table

| Reagent type<br>(species) or<br>resource | Designation | Source or<br>reference | Identifiers | Additional<br>information |
| --- | --- | --- | --- | --- |
| Gene<br>(*Arabidopsis thaliana*) | *TOPLESS, TPL* | GenBank | AT1G15750 |  |
| Gene<br>(*Arabidopsis thaliana*) | *TPR1* | GenBank | AT1G80490 |  |
| Gene<br>(*Arabidopsis thaliana*) | *TPR2* | GenBank | AT3G16830 |  |
| Gene<br>(*Arabidopsis thaliana*) | *TPR3* | GenBank | AT5G27030 |  |
| Gene<br>(*Arabidopsis thaliana*) | *TPR4* | GenBank | AT3G15880 |  |
| Gene<br>(*Arabidopsis thaliana*) | *ARF19* | GenBank | AT1G19220 |  |
| Gene<br>(*Arabidopsis thaliana*) | *IAA3/SHY2* | GenBank | AT1G04240 |  |
| Gene<br>(*Arabidopsis thaliana*) | *IAA14/SLR* | GenBank | AT4G14550 |  |

| Reagent type (species) or resource | Designation | Source or reference | Identifiers | Additional information |
| --- | --- | --- | --- | --- |
| Strain, strain background (*Saccharomyces cerevisiae*) | *AtARC*<sup>Sc</sup> strains | Pierre-Jerome et al., 2014; this study |  |  |
| Strain, strain background (*Escherichia coli*) | Dh5a |  |  | Cloning host |
| Strain, strain background (*Agrobacterium tumefaciens*) | GV3101 |  |  | Plant transformation |
| Genetic reagent (*Arabidopsis thaliana*) | *tpl-1* (SALK_097230) | ABRC | CS65909 | *mL* |
| Genetic reagent (*Arabidopsis thaliana*) | *tpr1-1 / mos10* | T-DNA induced deletion | This study; see Figure 1 - figure supplement 1C | *m1*; pSKI015 insertion, early stop at H455 |
| Genetic reagent (*Arabidopsis thaliana*) | *tpr2-3* | CRISPR insertion | This study; see Figure 1 - figure supplement 1D | *m2*; premature stop in CRA domain |
| Genetic reagent (*Arabidopsis thaliana*) | *tpr2-2* (SALK_079848C) | ABRC |  | *m2* |
| Genetic reagent (*Arabidopsis thaliana*) | *tpr3-1* (SALK_029936) | ABRC |  | *m3* |
| Genetic reagent (*Arabidopsis thaliana*) | *tpr4-1* (SALK_150008) | ABRC |  | *m4* |
| Genetic reagent (*Arabidopsis thaliana*) | J0121 enhancer trap line | Laplaze et al., 2005; Gala et al., 2021 |  | GAL4-VP16 expression in XPP cells |
| Genetic reagent (*Arabidopsis thaliana*) | pUBQ10::YFP-TPL iEraser | This study |  | See Figure 4B |
| Biological sample (*Botrytis cinerea*) |  |  |  | Pathogen challenge |
| Antibody | Anti-HA (Rat Monoclonal) |  |  |  |
| Antibody | Anti-Flag (Rat Monoclonal) |  |  |  |
| Recombinant DNA reagent | pUBQ10::YFP-TPL iEraser construct | This study |  |  |
| Recombinant DNA reagent | *AtARC*<sup>Sc</sup> Helix 8 variant constructs | This study |  |  |
| Recombinant DNA reagent | pAHP6::Effector constructs | This study |  |  |

| **Reagent type (species) or resource** | **Designation** | **Source or reference** | **Identifiers** | **Additional information** |
| --- | --- | --- | --- | --- |
| Recombinant DNA reagent | pARF19::mScarlet-P2A-RxL21 | This study |  |  |
| Chemical compound, drug | 2,4-Dichlorophenoxyacetic acid (2,4-D) | Fisher | ICN10155380 | Synthetic auxin for plant assays |
| Chemical compound, drug | Auxin (IAA) | PlantMedia.com | 705490 | 10 μM for yeast assays |
| Chemical compound, drug | Auxinole | MedChem Express | HY-111444 | 10 μM for plant assays |
| Software, algorithm | R | RStudio | rstudio.com |  |
| Software, algorithm | MEGA11 | Tamura et al., 2021 | megasoftware.net | Phylogenetic analysis |
| Software, algorithm | ImageJ/Fiji | Schneider et al., 2012 | imagej.nih.gov/ij/ |  |
| Software, algorithm | DESeq2 | Love et al., 2014 |  | Differential expression |
| Software, algorithm | tximport |  |  | RNA-seq read alignment |
| MacroPod |  |  |  | Confocal imaging |
| Other | BD special order cytometer | BD Biosciences |  | 514 nm laser; flow cytometry |

### Plant materials and growth conditions

All Arabidopsis thaliana lines used in this study are in the Columbia-0 (Col-0) accession. Seeds were surface-sterilized with 70% EtOH and 0.1% Triton-x and stratified at 4°C in the dark for 2 days. For root phenotyping experiments, seedlings were grown on 0.5× Linsmaier & Skoog (LS) medium supplemented with 0.8% agar, pH 5.7, constant light at 25°C. Plates were oriented vertically to allow root elongation along the agar surface. For rosette phenotyping and *B. cinerea* infection assays, plants were grown in neutral day conditions (12 hr light/12 hr dark) at 22°C and 60% humidity for 5 weeks, or until bolted for the bolting time experiment.

### Mutant alleles and genotyping

T-DNA insertion alleles for *tpl* (SALK_097230), *tpr2-2* (SALK_079848C), *tpr3* (SALK_029936), and *tpr4* (SALK_150008) were obtained from the Arabidopsis Biological Resource Center (ABRC). The *tpr1-1* (mos10) allele carries a pSKI015 T-DNA insertion that induces a deletion and early stop codon at H455 (Figure 1 -Figure Supplement 1C). Homozygosity of all T-DNA alleles was confirmed by PCR genotyping using gene-specific and T-DNA border primers (Supplementary File).

Higher-order mutant combinations (*mL124 and mL124,YL*) were generated by sequential crossing and confirmed by PCR genotyping at each generation.

### CRISPR-Cas9 mutagenesis of TPR2

The *tpr2-3* allele was generated using CRISPR-Cas9. Two guide RNAs targeting the Exon 4/5 junction of TPR2 (amino acid positions 152–192) was designed off of an approach based on a highly intron-optimized zCas9i gene and cloned into *At*-Fast with pDGE. The construct was introduced into *tpl tpr1 tpr4* by Agrobacterium-mediated floral dip transformation (Clough and Bent, 1998). T1 transformants were selected on hygromycin and Cas9-free T2 segregants carrying the desired insertion were identified by [Primers flanking insert site and Sanger sequencing]. The *tpr2-3* allele introduces premature stop codons within the CRA domain (Figure 1 - figure supplement 1D). Reduction of TPR2 expression in *tpr2-3* was confirmed by qPCR using four primer sets spanning positions upstream and downstream of the CRISPR target site (Figure1-figure supplement 1E).

### Integrase target cloning

The iEraser cassette was constructed by placing the YFP-TPL coding sequence under the constitutive pUBQ10 promoter, flanked by PhiC31 integrase recognition sites (attB and attP) in an invertible orientation, following the protocols established in ‘A Hot-Swappable Genetic Switch: Building an Inducible and Trackable Functional Assay for the Essential Gene MEDIATOR 21” (Watson et al., 2024). In the default orientation, YFP-TPL is expressed. Expression of PhiC31 integrase driven by the pARF19 promoter mediates site-specific inversion of the cassette in lateral root-associated cells, switching expression from YFP-TPL to mScarlet. The iEraser construct was introduced into *mL14* and *mL124* backgrounds by Agrobacterium-mediated transformation. Recombination events were confirmed by fluorescence imaging of mScarlet expression in lateral root primordia.

The iInducer cassette was constructed by placing the mScarlet-P2A-RxL21 coding sequence under the constitutive pUBQ10 promoter, flanked by PhiC31 integrase recognition sites (attB and attP) in an invertible orientation, following the protocols established in Watson et al., (2024). In the default orientation, the coding sequence is inverted relative to the promoter and mScarlet-P2A-RxL21 is not expressed. Expression of PhiC31 integrase driven by the pARF19 promoter mediates site-specific inversion of the cassette in lateral root-associated cells, placing the coding sequence in productive orientation and activating expression of mScarlet and RxL21 as separate polypeptides via P2A-mediated ribosomal skipping. This allows mScarlet to serve as a fluorescent reporter of recombination while RxL21 is delivered as an untagged EAR-containing effector capable of competing with Aux/IAA proteins for TPX binding. A parallel iInducer construct carrying a deletion of the LMLTL EAR motif (RxL21-ΔEAR) was generated as a non-repressive control. Both constructs were introduced into Col-0 by Agrobacterium-mediated transformation. Recombination events were confirmed by fluorescence imaging of mScarlet expression in lateral root primordia.

### Cloning and construct assembly

Construction of *At*ARC^Sc^ constructs was performed by Golden Gate cloning as described in Pierre-Jerome et al. (2014) and Leydon et al. (2021). In the fused configuration, the TPL N-terminal domain containing the Helix 8 variant of interest was directly fused to IAA3. In the unfused configuration, the Helix 8 variant-containing TPL N-terminal domain and IAA3 were expressed as separate proteins from individual promoters within the *At*ARC*^Sc^*. Helix 8 sequences corresponding to the L1 (TPL/TPR1), 23 (TPR2/TPR3), and 4 (TPR4) clades were synthesized using Q5 Site-Directed Mutagenesis (NEB, Cat #E0554S) and introduced into the appropriate *At*ARC^Sc^ backbone by Golden Gate assembly. All variant sequences were confirmed by Sanger sequencing (Genewiz/Azenta).

For effector competition assays, the HaRxL21 coding sequence was amplified and cloned into the *At*ARC^Sc^ unfused configuration as an additional expression cassette. An EAR-deletion variant of RxL21 (RxL21ΔEAR) was generated by PCR-mediated site-directed mutagenesis using Q5 Site-Directed Mutagenesis (NEB, Cat #E0554S).

For in planta effector expression apart from RxL21, pAHP6::Effector constructs were ordered as BsaI flanked gene fragments from Twist biosciences (San Francisco, USA) and assembled into the pGII backbone plasmid by Golden Gate cloning. The J0121 enhancer trap system driving GAL4-VP16 was used to express UAS::RxL21 constructs in xylem pole pericycle cells as described in Leydon et al. (2021). The pUAS::mScarlet-RxL21 construct was assembled using with mScarlet and RxL21 For UAS-driven constructs, the TPLN188-IAA14 coding sequence was amplified with primers containing engineered BsaI sites and introduced into the pGII backbone with the UAS promoter and RBSC terminator (Siligato et al., 2016) using Golden Gate cloning as described previously (Leydon et al., 2021).

### Yeast methods and flow cytometry

Standard yeast synthetic dropout (SDO) and yeast extract–peptone–dextrose plus adenine (YPAD) media were used. A standard lithium acetate protocol (Gietz and Woods, 2002) was used for transformations. All cultures were grown at 30°C with shaking at 250 rpm.

For *At*ARC*^Sc^* repression assays, yeast strains carrying the integrated circuit components were grown overnight in SDO medium, diluted to ∼100 events/µL as determined by flow cytometry, and grown for an additional 4–6 hr to reach mid-log phase before measurement. For auxin-responsive experiments, “*auxin need to check with alex*” (Auxin, 10 µM) was added at the indicated time. Fluorescence measurements were taken using a Becton Dickinson (BD) special order cytometer with a 514 nm laser exciting mVenus fluorescence with a 525 nm cutoff filter prior to photomultiplier tube collection. A total of 5,000–10,000 events were collected for each sample. Data were exported as FCS 3.0 files and processed using the flowCore R package and the FlowTime R package (Wright et al., 2019). Fluorescence values were normalized to cell size (FSC-A). Data from at least two independent biological and two independent technical replicates were combined and plotted using ggplot2 in R. 95% confidence intervals were calculated and displayed for each construct.

### Root growth and auxin sensitivity assays

For dose-response curves to auxin, primary root length was measured from scanned images of seedlings grown vertically on 0.5× LS plates containing 2,4-D at concentrations of 0, 2, 6.5, 20, and 60 nM. For single-concentration treatments, seedlings were grown vertically on 0.5× LS plates supplemented with either 6.5 nM 2,4-D or 0.5 μM auxinole (a competitive inhibitor of the TIR1/AFB auxin co-receptors; (Hayashi et al., 2012), with mock (solvent-only) plates used as controls. Auxinole was dissolved in DMSO and added to autoclaved media cooled to ∼55 °C immediately before pouring; final DMSO concentration was kept constant across auxinole and mock plates. Seedlings were imaged at 4 days after transplanting using an Epson V800 flatbed scanner, and root lengths were measured using ImageJ/Fiji ((Schneider et al., 2012). Dose-response curves were fitted using a four-parameter log-logistic model in R (drc package;(Ritz et al., 2015). Statistical comparisons between genotypes and treatments were performed using ANOVA followed by Tukey’s HSD post-hoc test.

### Lateral root phenotyping

Lateral root density was quantified by counting emerged lateral roots on seedlings grown vertically on 0.5xLS plates at 14 days after germination, unless otherwise indicated. Root length was measured from scanned plate images, and lateral root density was calculated as the number of emerged lateral roots per centimeter of primary root. For time-course analyses (Figure 5J,K), lateral root emergence was scored daily between 10 and 14 days after germination and were measured using ImageJ/Fiji (Schneider et al., 2012).

### Gravistimulation

For gravistimulation-induced lateral root initiation experiments, seedlings were grown vertically for 3 days on 0.5x LS plates and then rotated 90° to induce a gravitropic bend. The proportion of seedlings initiating lateral roots at the bend site was scored at 1 through 6 days after rotation.

### Root hair analysis

Root hair length and density were assessed from DIC micrographs of the root hair differentiation zone. Seedlings were imaged using a Leica DMI3000B at objective level 10x. Root hair length was measured using ImageJ/Fiji

### Fluorescent microscopy

Fluorescent imaging of YFP-TPL and mScarlet reporters in iEraser lines was performed on a Leica DMI3000B and imaged with Leica DFC 345 FX. YFP was excited at 514 nm; mScarlet was excited at 561 nm. For GFP imaging in J0121 enhancer trap lines, excitation was at 488 nm. Images were processed in ImageJ/Fiji.

### Floral phenotyping

Petal number was scored on the first 150 flowers per genotype. Flowers were examined under a dissecting microscope Leica S8AP0 and imaged with Leica DMC 2900. Floral organ identity defects were documented photographically and categorized qualitatively. Flowers were also examined under a MacroPod PRO, MP-PRO-3D-5DSR, to generate high resolution photos. Full resolution files of MacroPod Flower images located in Supplementary File.

### RNA extraction and quantitative RT-PCR

0.5x MS plate grown 10-day-old seedling RNAs were extracted using Qiagen RNeasy. cDNA synthesis was performed using SuperScript IV (Invitrogen) with oligo(dT) oligos. cDNA was diluted 1/10 and 2 μl used for final qPCR. SYBR Green Master Mix (Applied Biosystem) was used for qPCR with a QuantStudio 3 machine (Thermo Fisher). Gene expression was normalised using UBQ11. The standard 2-ΔΔCt method was used to quantify mRNA levels. Primer sequences are listed in Supplementary File.

### Chromatin immunoprecipitation (ChIP)

ChIP assays were performed as described previously (Xing et al., 2013) with the following modifications: ½ MS plate grown seedlings (10-day old) were used. Sonication was performed using a Bandolin Sonopuls HD 2070 sonicator at 40 % duty cycle and 20 % power with a MS73 probe (4 cycles). The antibody used was anti-HA (Abcam ab9110). DNA was purified using the Qiagen Qiaquick PCR purification kit eluting in a final 200μl volume. The TPL::TPL-HA and TPR1::TPR1-HA transgenic lines used were described previously (Szemenyei et al., 2008; Zhu et al., 2010).

### Infection assays and transcriptomic analysis

The *B. cinerea* pepper isolate (Denby et al., 2004) was grown in the dark on apricot halves at 25°C for two weeks before spore collection. 10 µL of a spore solution (1.5 x 10^6^ spores/mL in half-strength grape juice) was inoculated onto detached Arabidopsis leaves, which were placed on 0.8% agar and incubated in a growth chamber in neutral days and 90% humidity. For the RNA-seq experiment, samples of mock and inoculated leaves (three independent pools of three leaves for each genotype/condition) were taken at 16 and 20 hpi and flash-frozen in liquid nitrogen. For disease susceptibility assays, lesion measurements were taken at 72 hpi and analyzed using Fiji image analysis software (Schindelin et al., 2012).

For RNA-seq,total RNA was extracted using the RNeasy Plant Mini Kit (Qiagen). Library preparation was performed using the Novogene NGS RNA Library Prep Set (PT042) and sequenced on the NovaSeq X Plus platform with paired-end 150 bp reads, generating approximately 6 Gb of data per sample.

Reads were quality-trimmed using fastp (Chen et al., 2018) and aligned to a combined AtRTD3 and *B. cinerea* ASM83294v1 reference transcriptome using Salmon (Patro et al., 2017; Zhang et al., 2022). Read counts were generated using tximport (Soneson et al., 2016) and differential expression analysis was performed using DESeq2 (Love et al., 2014) with an adjusted p-value threshold of 0.01 and a log2 fold-change cutoff of ±0.5. Venn diagrams were generated using ggplot. Hierarchical clustering based on Euclidean distances and heatmap visualization were performed in R using pheatmap (Kolde, 2025). Gene Ontology enrichment analysis was performed using ShinyGO v0.85.1(Ge et al., 2020).

### Leaf and bolting phenotyping

For measuring leaf number and area, the leaves of three plants per genotype were removed after they had been grown in neutral days for 5 weeks. Leaf area was measured using Fiji image analysis software. To check for statistical differences in leaf area between genotypes, a linear mixed-effects model was fitted using the nlme package (Pinheiro et al., 2026). Pairwise comparisons were calculated with the emmeans package (Lenth and Piaskowski, 2026).

For the bolting time series, 20 plants for each genotype were observed for the time they took to reach bolting. Bolting was defined by the main stem and bud together reaching a length of 1 cm. To check for statistical differences in bolting time, a log rank test was performed using the survival package (Therneau, 2026).

### Phylogenetic analysis

Protein sequences of the TPX family (TPL, TPR1–4) were aligned using ClustalW within MEGA11 (Tamura et al., 2021). Evolutionary relationships were inferred using the Maximum Likelihood method and JTT matrix-based model. Bootstrap support was assessed with 100 replicates. Helix 8 sequences were extracted from the full-length alignment and analyzed separately to generate the clade-specific phylogeny shown in Figure 3B.

### Arabidopsis transformation

Stable transgenic Arabidopsis lines were generated by *Agrobacterium tumefaciens* (strain GV3101)-mediated floral dip transformation, T1 transformants were selected on LS with Hygromycin for selection. For constructs driven by the J0121 enhancer trap, T1 seeds were screened for fluorescence or resistance. Single-insertion lines were identified by segregation analysis in the T2 generation, and homozygous T2 lines were used for phenotypic analysis unless otherwise stated. For pAHP6::Effector lines, T2 populations were used as indicated in Figure 5F.

### Statistical analysis

Statistical analyses were performed in R (version 2023.06.0+421). For comparisons between two groups, Student’s t-test was used. For comparisons among multiple genotypes, one-way ANOVA followed by Tukey’s HSD post-hoc test was used unless otherwise specified. Sample sizes and statistical tests are indicated in the corresponding figure legends. Throughout, p < 0.05 was considered statistically significant unless stated otherwise. Exact p-values are reported where applicable.

## Data and code availability

All raw sequencing data have been deposited at <u>DOI: 10.5061/dryad.2rbnzs84g</u>. Raw RNAseq data is deposited at NCBI under Project <u>PRJNA1465640</u>. Custom analysis scripts are available at https://github.com/achillobator/TPX. Seed stocks for novel mutant alleles and transgenic lines described in this study will be deposited at the ABRC.

## Acknowledgements

The work in the Nemhauser and Denby Labs is supported by the National Institutes of Health (R35-GM148135), and the Benjamin D. Hall Endowed Chair in Basic Life Science. BD was additionally supported by Tunnicliffe and WRF-Hall Fellowships. MP is funded by the BBSRC White Rose Doctoral Training Partnership in Mechanistic Biology (BB/T007222/1) and FV by University of York funds awarded to KD.

## Additional Information

### Author Contributions

**Benjamin L.R. Downing**

Department of Biology, University of Washington, Seattle, United States

<u>Contribution:</u> Conceptualization, Data curation, Formal analysis, Validation, Investigation, Visualization, Methodology, Writing - original draft, Project administration, Writing - review and editing

<u>Competing interests:</u> No competing interests declared

**Maria Pattichis**

Center for Novel Agricultural Products (CNAP), Department of Biology, University of York, York, United Kingdom

<u>Contribution:</u> Conceptualization, Data curation, Formal analysis, Validation, Investigation, Visualization, Methodology, Writing - original draft, Project administration, Writing - review and editing

<u>Competing interests:</u> No competing interests declared

**Fabian E. Vaistij**

Center for Novel Agricultural Products (CNAP), Department of Biology, University of York, York, United Kingdom

<u>Contribution:</u> Conceptualization, Data curation, Formal analysis, Validation, Investigation, Visualization, Methodology, Writing - original draft, Project administration, Writing - review and editing

<u>Competing interests:</u> No competing interests declared

**Mohamed Farawila**

Department of Biology, University of Washington, Seattle, United States <u>Contribution:</u> Investigation, Visualization, Writing - review and editing <u>Competing interests:</u> No competing interests declared

**Edward Ghannam**

Department of Biology, University of Washington, Seattle, United States <u>Contribution:</u> Investigation, Visualization, Writing - review and editing <u>Competing interests:</u> No competing interests declared

**Linda Nguyen**

Department of Biology, University of Washington, Seattle, United States <u>Contribution:</u> Investigation, Visualization, Writing - review and editing <u>Competing interests:</u> No competing interests declared

**Katherine Denby**

Center for Novel Agricultural Products (CNAP), Department of Biology, University of York, York, United Kingdom

<u>Contribution:</u> Conceptualization, Resources, Supervision, Funding Acquisition, Project administration, Writing - review and editing

For Correspondence

<u>Competing interests:</u> No competing interests declared

**Alexander Leydon**

Department of Biology, University of Washington, Seattle, United States

<u>Contribution:</u> Conceptualization, Data curation, Formal analysis, Validation, Investigation, Visualization, Methodology, Writing - original draft, Project administration, Writing - review and editing

<u>Competing interests:</u> No competing interests declared

**Jennifer Nemhauser**

Department of Biology, University of Washington, Seattle, United States

<u>Contribution:</u> Conceptualization, Resources, Supervision, Funding Acquisition, Project administration, Writing - review and editing

For Correspondence

<u>Competing interests:</u> No competing interests declared

## Funding

**National Institutes of Health (R35-GM148135)**

- Benjamin L.R. Downing
- Mohamed Farawila
- Edward Ghannam
- Linda Nguyen
- Alexander Leydon

**BBSRC White Rose Doctoral Training Partnership in Mechanistic Biology (BB/T007222/1)**

- Maria Pattichis

## Additional Files

**Table_S1:** *Botrytis cinerea* effectors

**Table_S2:** Gene expression values from profiling of mock- (M) and *Botrytis cinerea* (B) inoculated Arabidopsis leaves from wildtype (WT), *tpl tpr1* (L1), *tpl tpr1 tpr4* (L14) and *tpr2 tpr3* (T23) mutants at two time points after inoculation (16 and 20 hours) for three biological replicates per sample. Genes differentially expressed in *tpx* mutants compared to wildtype in the mock-inoculated samples. Genes differentially expressed in *B. cinerea* inoculated samples compared to mock inoculated.

**Table_S3:** Primer List

